# Large-Scale Analysis of Cell Death Phenotypic Heterogeneity

**DOI:** 10.1101/2020.02.28.970079

**Authors:** Zintis Inde, Giovanni C. Forcina, Kyle Denton, Scott J. Dixon

## Abstract

Individual cancer cells within a population can exhibit substantial phenotypic heterogeneity such that exposure to a lethal agent will kill only a fraction of cells at a given time. Whether common molecular mechanisms govern this fractional killing in response to different lethal stimuli is poorly understood. In part, this is because it is difficult to compare fractional killing between conditions using existing approaches. Here, we show that fractional killing can be quantified and compared for hundreds of populations in parallel using high-throughput time-lapse imaging. We find that fractional killing is highly variable between lethal agents and between cell lines. At the molecular level, we find that the antiapoptotic protein MCL1 is an important determinant of fractional killing in response to mitogen activated protein kinase (MAPK) pathway inhibitors but not other lethal stimuli. These studies lay the foundation for the large-scale, quantitative analysis of cell death phenotypic heterogeneity.

## INTRODUCTION

Individual cells within a population often behave differently even when exposed to the same environment. This heterogeneity is apparent in both prokaryotic and eukaryotic cells, and affects a wide range of phenotypes including morphology, chemotaxis, and the response to lethal drug treatment (Bigger, 1944; Santos et al., 2019; Sharma et al., 2010; Spudich and Koshland, 1976; Wu et al., 2020). Such differences can arise in a non-genetic manner due to random fluctuations within molecular networks impacting gene or protein expression, epigenetic states, metabolism and other factors (Niepel et al., 2009). It is of great interest to understand the nature of this heterogeneity at the cellular and molecular levels and how it contributes to specific biological outcomes.

Phenotypic heterogeneity is apparent in the response of cancer cells to lethal drugs. At a simple level, lethal drugs can generally be titrated such that only a fraction of the total number of cells within a population are killed at a given dose and time (Figure 1A) (Fallahi-Sichani et al., 2013). This heterogeneity may be related to fractional killing (FK), a clinical phenomenon where a constant fraction of tumor cells are killed in response to each cycle of drug administration rather than a constant number (Berenbaum, 1972; Skipper, 1965). FK provides a useful conceptual paradigm within which to examine cell death phenotypic heterogeneity in vitro. Here, FK has been linked to differences between cells in the expression or activity of several pro-apoptotic and signaling molecules, including caspases, p53, and the c-Jun N-terminal kinase (JNK) pathway (Miura et al., 2018; Paek et al., 2016; Roux et al., 2015; Spencer et al., 2009). How FK varies between lethal conditions, and whether FK in response to different lethal stimuli is governed by a core molecular mechanism or is stimuli-specific, is largely unclear. In part, this is due to the low-throughput nature of existing methods to assess FK.

**Figure 1.**
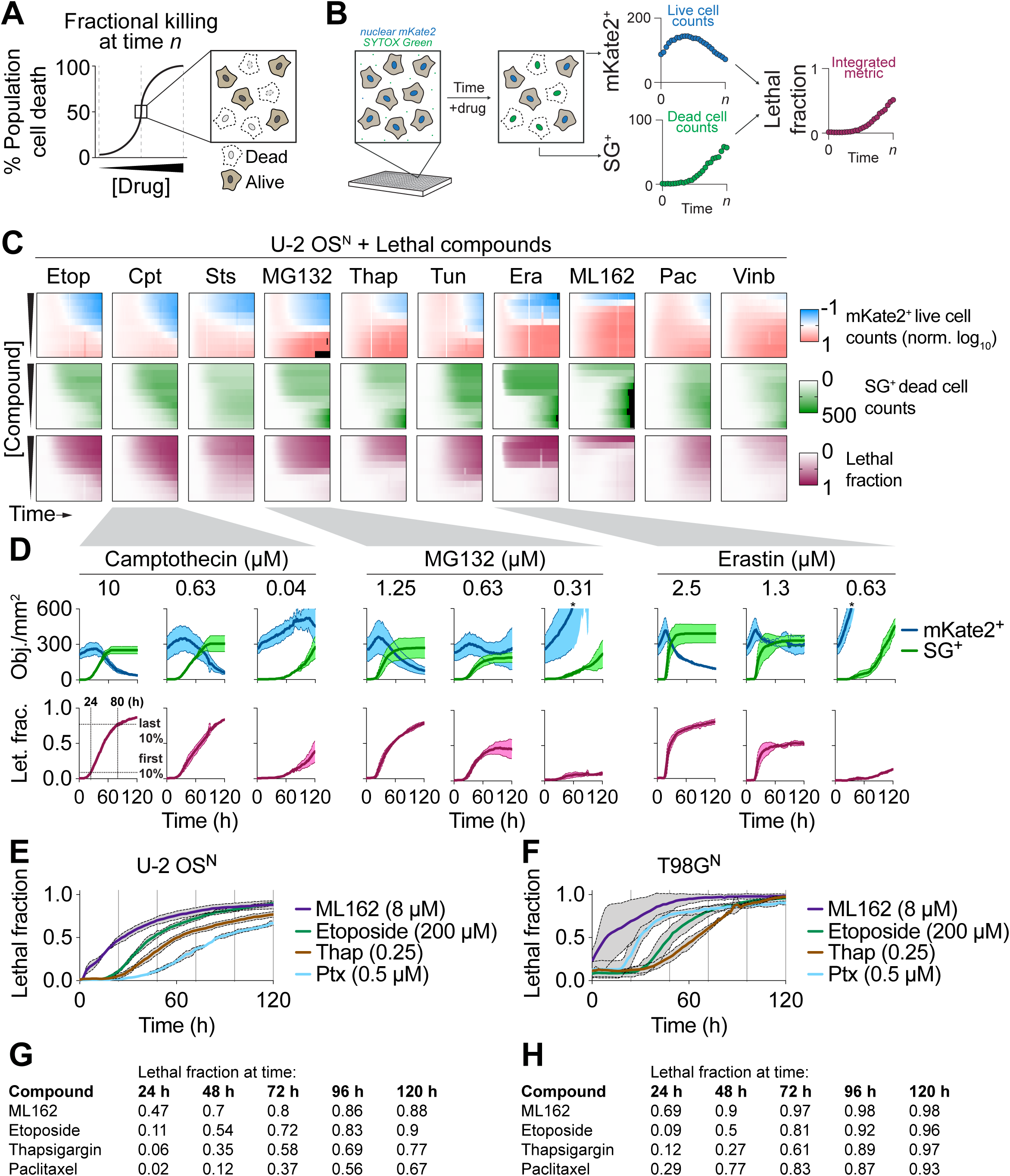
Systematic investigation of fractional killing. (A) At a given drug dose and time, a fraction of cells within the population will be killed while others survive. (B) Overview of cell death analysis using time lapse imaging of population live (nuclear mKate2) and dead (SYTOX Green) cell over time. These data can be integrated into a single metric, the lethal fraction, representing overall population cell death. (C) Heatmap summaries of mKate2 positive (mKate2^+^) live cell counts, SYTOX Green positive (SG^+^) dead cell counts, and integrated lethal fraction scores as a function of time (x-axis) and concentration (y-axis) for ten compounds in U-2 OS^N^ cells. For the mKate2^+^ heatmaps, live cell counts were normalized to 1 at time = 0 and expressed on a log_10_ scale. Etop: etoposide, Cpt: camptothecin, Sts: staurosporine, Thap: thapsigargin, Tun: tunicamycin, Era: erastin, Pac, paclitaxel, Vinb: vinblastine. (D) Isolated live cell counts, dead cell counts (mKate2^+^ or SG^+^ objects/mm^2^ imaged area; Obj./mm^2^) and corresponding lethal fraction (Let. frac.) scores over time, extracted from select conditions in C. The stars (*) indicate where live cell counts increase beyond the range of the y-axis scale. (E, F). Lethal fraction scores over time for U-2 OS^N^ (E) and T98G^N^ (F) cells exposed to four lethal compounds. (G, H). Extracted mean lethal fractions at select timepoints from the data presented in E and F. Data are from three independent experiments and represent the mean (C, G, H) or mean ± SD (D-F).

Here, we examine phenotypic heterogeneity within the framework of drug-induced cancer cell FK. In particular, we sought to determine whether the nature of FK varies between lethal conditions and is regulated by common or context-specific mechanisms. We find that it is possible to systematically quantify FK under many conditions in parallel using high-throughput population-level time-lapse imaging and associated mathematical modelling. We identify the anti-apoptotic protein MCL1 specifically as one important regulator of FK in response to inhibitors of the mitogen activated protein kinase (MAPK) cascade, but not other lethal stimuli. Our results demonstrate that it is possible to analyze FK on a large-scale basis and suggest that this phenomenon is governed by diverse molecular mechanisms.

## RESULTS

### Traditional cell viability assays do not report effectively on fractional killing

We sought to determine whether shared molecular mechanisms regulate FK in response to different lethal stimuli and across genetic backgrounds. We also reasoned that it would be important to disentangle the potential effect of drug dose on FK (e.g. intermediate drug doses may yield FK simply due to variable target engagement between cells in the population (Mateus et al., 2017)). Thus, for this analysis, the ideal approach would enable us to assess FK accurately for many different drug treatment conditions in parallel. At the beginning of our studies, the best approach was unclear. Population cell death is typically inferred in large-scale studies using metabolism-based cell viability assays (Kepp et al., 2011). These assays are simple, affordable, and highly accessible. To test if these methods would be suitable, we examined cell viability in two genetically distinct cancer cell lines, U-2 OS^N^ osteosarcoma and T98G^N^ glioblastoma, using two common metabolism-based assays, CellTiter-Glo (CTG) and PrestoBlue (PB). As test lethal perturbations, both cell lines were treated with a ten-point, two-fold dose-response series of three mechanistically-distinct lethal compounds (etoposide, bortezomib and vinblastine) for 72 h. U-2 OS^N^ and T98G^N^ cells express the live cell marker nuclear-localized mKate2 (denoted by the superscript ‘N’, (Forcina et al., 2017)). Crucially, this enabled us to benchmark CTG and PB measurements against live cell counts from these same populations and examine how CTG and PB measures compared to actual cell counts.

As expected, CTG and PB measurements and live cell counts declined with increasing concentrations of all three compounds in both cell lines (Figure S1A-D, and *not shown*). However, changes in CTG and PB measurements frequently did not correlate linearly with live cell counts. For example, in U-2 OS^N^ cells, treatment with 400 nM or 1.6 µM etoposide resulted in CTG and PB measurements that were within error of one another, despite differing by 35% in live cell counts (Figure S1A,C). Similar discrepancies were likewise observed in T98G^N^ cells treated with bortezomib or vinblastine (Figure S1B,D). We also observed that CTG and PB measurements were not constant on a per cell basis, being substantially increased at higher compound concentrations, possibly due to compound effects on mitochondrial function (Chan et al., 2013) (Figure S1E,F). Per cell CTG and PB values also varied considerably between U-2 OS^N^ and T98G^N^ cell lines (Figure S1C,E). Collectively, these results suggested that metabolism-based measures of cell viability would not be suitable for a comparative analysis of heterogeneous cell death between populations.

### Analyzing fractional killing using high-throughput time-lapse microscopy

Our results prompted us to consider alternative methods for evaluating FK. In particular, we assessed the utility of high-throughput time-lapse imaging. This approach is less accessible than traditional cell viability assays, but can enable both live (mKate2^+^) and dead (SYTOX Green-positive) cells to be counted in many populations in parallel over time, as we have recently demonstrated (Figure 1B) (Forcina et al., 2017). Accordingly, we hypothesized that high-throughput time-lapse imaging of live and/or dead cells would enable effective quantification of FK across conditions. To begin testing this hypothesis, we examined populations of U-2 OS^N^ and T98G^N^ cells treated with a panel of ten different lethal compounds, each tested in 10-point, two-fold dose-response series. For all 200 conditions, live and dead cells were counted every 2 h for a total of 120 h and the resultant dataset visualized using heatmaps (Figure 1C, S1G).

All compounds reduced live cell counts and increased dead cell counts at one or more tested concentrations in both cell lines (Figure 1D, S1G). At high drug concentrations, we typically observed a decrease in live cells by the end of the observation period that equaled the increase in dead cells. However, at intermediate lethal doses, live or dead cell counts alone did not accurately reflect total population cell death over time. For example, in U-2 OS^N^ cells treated with an intermediate dose of camptothecin (0.04 µM), MG132 (0.63 µM) or erastin (1.3 µM), mKate2^+^ live cell counts at 120 h were similar to those observed at the start of the experiment, despite extensive cell death within the population as indicated by increased SG^+^ dead cell counts (Figure 1D). Similar observations were made in T98G^N^ cells treated with intermediate concentrations of MG132 or thapsigargin (Figure S1H). This effect was best explained by the fact that most lethal treatments did not start killing cells immediately upon addition. Rather, populations typically increased in cell number for several hours prior to the onset of cell death, expanding the initial size of the population subsequently available to die (Figure 1C,D, S1G,H). We therefore concluded that to assess FK it was essential to track both live and dead cells, as done in low-throughput experimental designs (Miura et al., 2018; Paek et al., 2016; Spencer et al., 2009).

Counts of live and dead cells alone do not provide a uniform metric for FK that is easy to compare across conditions. We previously described how counts of live and dead cells over time could be integrated into a single metric, the lethal fraction, which summarizes overall population cell death (Forcina et al., 2017) (Figure 1B). We hypothesized that lethal fraction scores would be a useful metric for FK within any given population. In support of this hypothesis, lethal fraction scores correctly accounted for the partial population cell death observed at intermediate compound doses noted above (Figure 1C,D, S1G,H). Interestingly, examination of lethal fraction scores over time made clear that FK is inherently a time dependent phenomenon. For example, in U-2 OS^N^ cells treated with 10 μM camptothecin, the first 10% of cells died by 24 h while the last 10% of cells did not die until after 80 h of treatment (Figure 1D). Similar variability in the span of time between the first and last cell deaths within the population were apparent for all lethal compounds examined, and in some cases (e.g. paclitaxel) this varied considerably between cell lines (Figure 1E-H). We conclude that high-throughput time-lapse imaging and lethal fraction scoring comprise a useful approach to assess FK on a common scale that can be compared across conditions.

### Fractional killing kinetics vary between lethal conditions

Lethal fraction scores provided a useful means to visualize different patterns of FK over time in response to different lethal treatment. However, further abstraction of these data was necessary to simplify the process of comparing FK between conditions. Recently we showed that lethal fraction scores over time can be fitted with a lag exponential death (LED) model that allows for the extraction of two key parameter values describing cell death within the population: D_O_ and D_R_ (Figure 2A) (Forcina et al., 2017). Previously we focused on D_O_, which reflects the time when the first cells in a population begin to die. By contrast, D_R_ captures the maximal rate of cell death within the population (i.e. the maximum slope of the lethal fraction curve) and is represented here as the rate of change in lethal fraction (expressed as a percentage of dead cells) over time (% h^-1^). D_R_ reflects the degree of heterogeneity in the initiation of cell death *between* cells within the population: high D_R_ values are observed when cells within the population die all around the same time, while low D_R_ values indicate that cells within the population are dying at very different times (Figure 2B). We therefore hypothesized that the D_R_ parameter value would provide a powerful means to summarize FK within a population and compare these values across conditions.

**Figure 2.**
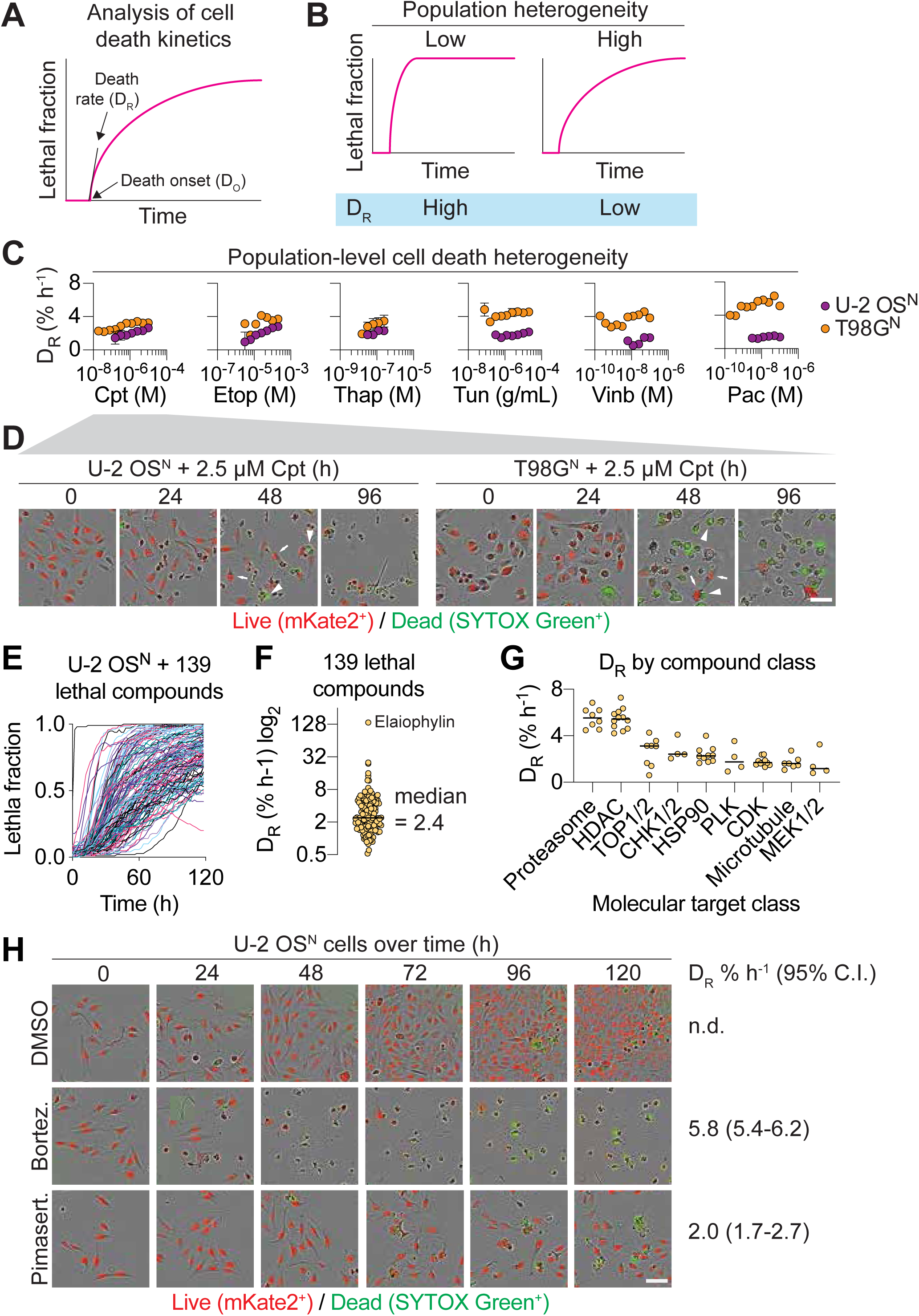
Quantification of time-dependent fractional killing. (A) Overview of cell death kinetic analysis using time-lapse imaging and curve fitting. Lethal fraction curves can be fit with a lag exponential death (LED) model that yields two key parameter values, D_O_ and D_R_, which reflect the timing of cell death onset and the maximum rate of cell death within the population, respectively (Forcina et al., 2017). (B) Treatments that trigger cell death with low between-cell heterogeneity will have high D_R_ values, and vice versa, which can impact the interpretation of FK at any given early timepoint. (C) Extracted D_R_ parameter values for LED curves fit to lethal fraction scores for a sub-set of the lethal compounds shown in Figure 1D and S1G. Kinetic parameter values were only computed for compound concentrations where the lethal fraction at 120 h exceeded 0.5. Results are mean ± SD from three independent experiments. (D) Representative population images over time for camptothecin (Cpt)-treated cells. Heterogeneity is especially apparent at the 48 h timepoint, where arrows indicate example live cells and arrowheads indicate example dead cells. Scale bar = 75 µm. (E) Traces of lethal fraction scores over time for 140 lethal compounds identified previously. (F) Extracted D_R_ values for 139 lethal compounds from E where it was possible to compute this parameter value. (G) D_R_ values for compounds in F broken down by compound class. (H) Representative population images for select conditions from G to illustrate heterogeneity in cell death initiation throughout the population. Scale bar = 75 µm.

To test this hypothesis, we examined how D_R_ varied for a subset of lethal conditions examined in U-2 OS^N^ and T98G^N^ cells. In principle, a condition where cell death is initiated at the same time in all cells within the population could have D_R_ equal to ∼100% h^-1^. Strikingly, while D_R_ varied between lethal compounds, it rarely exceeded 6% h^-1^ (Figure 2C). In fact, in many cases D_R_ was below 3% h^-1^, even at the highest tested compound concentrations. These D_R_ values suggest that cells within the population execute cell death at very different times, a phenotype we confirmed by visual inspection (Figure 2D, S2A,B). D_R_ captures this cellular heterogeneity and illustrates the degree to which FK within a population is fundamentally time dependent. Notably, this time dependence could not easily have been captured by endpoint measurements of CellTiter-Glo and PrestoBlue metrics.

To further test our hypothesis, we examined how FK varied between 140 mechanistically-diverse lethal compounds tested in U-2 OS osteosarcoma cells at a single dose of 5 µM (Figure 2E) (Forcina et al., 2017). 139 of 140 compounds yielded lethal fraction scores over time (120 h of observation) where it was possible to determine D_R_ values. Across all compounds, median D_R_ was 2.4% h^-1^ (1.6 - 4.6% h^-1^ interquartile range) (Figure 2F), consistent with analysis of a more limited set of lethal compounds above. Most compounds in this set have a specific molecular target, allowing us to ask whether FK varied by compound target. Consistent with our previous analysis (Forcina et al., 2017), D_R_ varied by compound class, with proteasome inhibitors and HDAC inhibitors triggering relatively fast FK (median D_R_ > 5% h^-1^), and inhibitors of cyclin-dependent kinases (CDK), microtubule function, and especially mitogen-activated protein kinase kinase 1 and 2 (MEK1/2) causing relatively ‘slow’ FK (median D_R_ < 2% h^-1^) (Figure 2G,H). Thus, the rate of FK over time can vary substantially between lethal compound classes.

### MEK1/2 inhibitors trigger slow FK not involving many known mechanisms

MEK1/2 inhibitors (MEKis) block the function of the important RAS/MAPK signal transduction pathway (Figure 3A). The slow kinetics of MEKi-induced FK were intriguing. We sought to identify molecular mechanisms that modulate this process and test whether these mechanisms were generalizable to other lethal compounds. MEKis are being investigated for use in the treatment of various cancers, including lung (Kim and Giaccone, 2018; Robert et al., 2019). We therefore pursued our follow-on studies using non-small cell lung carcinoma (NSCLC) models. Crucially, the structurally-distinct MEKis pimasertib (Pim) and trametinib (Tram) triggered slow FK (D_R_ < 3% h^-1^) across a range of lethal concentrations in three different KRAS-mutant NSCLC cell lines, A549, Calu-6 and H2291, consistent with a broadly generalizable FK phenotype (Figure 3B,C, S3A).

**Figure 3.**
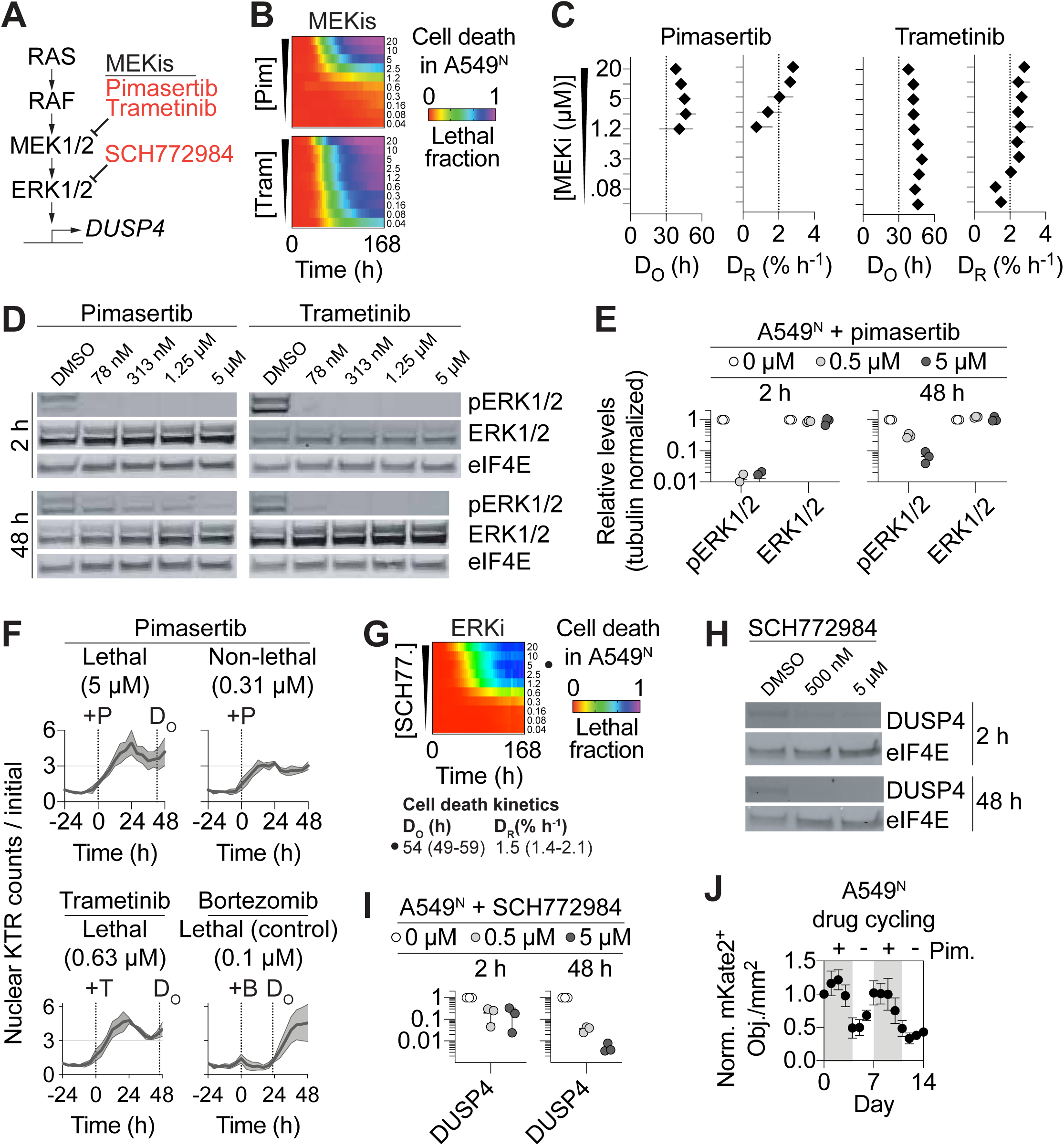
MEKis trigger cell death with high between-cell variability. (A) Outline of key RAS/MAPK pathway components highlighting inhibitors (orange) of MEK1/2 (i.e. MEKis) and ERK1/2. (B) Cell death in A549^N^ cells summarized over time and MEKi concentration. (C) Cell death kinetic parameters for the data in B summarized over compound concentration. Note: parameters can only be computed with confidence for compound concentrations with lethal fraction > ∼0.2. (D) Western blot for phosphorylated and total ERK1/2 in A549^N^ cells. (E) Quantification of phospho-ERK1/2 and total ERK1/2 at different concentrations of pimasertib at two timepoints. (F) Data for ERK kinase translocation reporter (KTR) in A549^N^ cells in response to MEKi treatment over time. ERK KTR response to an unrelated lethal compound, bortezomib, is shown as a control. Death onset for lethal conditions (from B,C for MEKis and not shown for Btz) is shown by a dashed line (D_O_) (G) Cell death in A549^N^ cells summarized over time and concentration for the ERK inhibitor (ERKi) SCH772984. Cell death kinetic parameter values are shown for cells treated with 5 µM SCH772984. (H) Expression of the MAPK pathway target DUSP4 in response to SCH772984. (I) Quantification of experiment outlined in H, for three individual experiments. (J) Normalized live cell (mKate2^+^ objects) counts within the same population of cells over two cycles of pimasertib (Pim, 5 µM, grey shaded area) addition with an intervening period of regrowth in the absence of drug (white area). All data are from at least three independent experiments, and represented as the mean (B,G), mean ± 95% C.I. (C), or mean ± SD (E,F,I,J).

A trivial explanation for FK is variable inhibition of the drug target. However, phosphorylation of the MEK1/2 substrates ERK1/2 was fully inhibited by Pim in as little as 2 h, long before the initiation of cell death, with a lethal concentration (i.e. 5 µM) resulting in sustained inhibition of ERK1/2 phosphorylation over 48 h (Figure 3D,E). Using the single-cell ERK kinase translocation reporter (KTR) (Regot et al., 2014), we found that lethal MEKi treatment resulted in greater reporter nuclear translocation before the onset of cell death than non-lethal MEKi treatment or lethal control (i.e. bortezomib) treatment (Figure 3F). Thus, differential inhibition of ERK1/2 phosphorylation did not explain slow FK in response to MEKis.

We next sought to identify proteins or processes downstream of MEK1/2 that could mediate slow FK. Similar to MEKis, direct inhibition of ERK1/2 using the small molecule ERK1/2 inhibitor (ERKi) SCH772984 resulted in slow FK in both A549^N^ and Calu-6^N^ cells (Figure 3G, S3B). We confirmed that SCH772984 effectively reduced the expression of DUSP4, a target of the MAPK pathway, at both 2 h and 48 h (Cagnol and Rivard, 2013) (Figure 3H,I). The similar MEK1/2i and ERKi phenotypes argued against the existence of an inherently drug resistant (i.e. mutant) sub-population. Indeed, Pim triggered FK with similar kinetics in both a drug-naïve cells and cells that survived an initial round of MEKi treatment (Figure 3J). Rather, we hypothesized that slow FK was due to non-genetic molecular heterogeneity downstream or in parallel to ERK1/2. Several known candidate mechanisms of FK (Inde and Dixon, 2018), including p53 stabilization, cell cycle phase, glycolytic or oxidative metabolism, and reactive oxygen species (ROS) accumulation, did not contribute to MEKi-induced FK (Figure S3C-J). Thus, the molecular heterogeneity contributing to FK in response to MAPK pathway inhibition appeared to reside elsewhere in the cellular network.

### Variable MCL1 expression contributes to slow FK

How inhibition of the MAPK pathway induced FK with such slow kinetics remained unclear. Given our ability to quantify FK for multiple independent conditions in parallel, we conducted a series of mechanistic investigations to explore this question. Testing the role of protein synthesis, we found that MEKi-induced cell death was potently suppressed by co-treatment with cycloheximide, suggesting that the translation of certain gene products was necessary for cell death (Figure S4A). Using RNA sequencing (RNA-Seq) we found a set of 555 genes significantly upregulated by lethal (i.e. 5 µM) Pim treatment in both A549^N^ and Calu-6^N^ cells that was enriched for regulators of apoptosis (Figure 4A,B). Indeed, cell death induced by MEKis or the positive control bortezomib was suppressed by co-treatment with the pan-caspase inhibitor Q-VD-OPh (Figure S4B). Inhibition of the MAPK pathway can induce apoptosis, and variable expression of apoptotic proteins is linked to FK in other contexts (Boucher et al., 2000; Hata et al., 2014; Roux et al., 2015; Spencer et al., 2009; Zhao and Adjei, 2014). Accordingly, we focused on how the apoptosis pathway impacted slow FK in response to MEKis.

**Figure 4.**
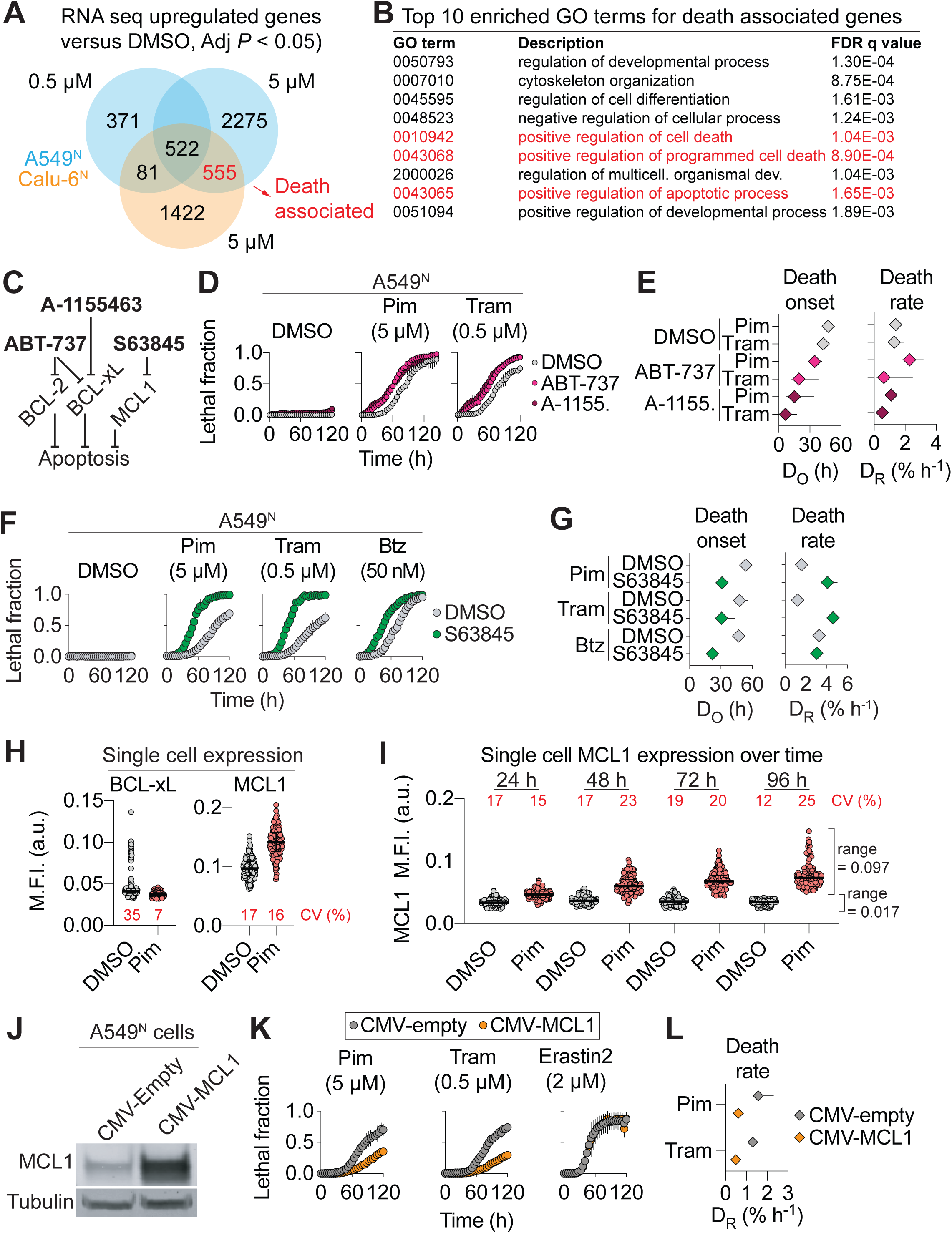
Variable Mcl-1 expression correlates with MEKi-induced FK. (A) Summary of RNA sequencing analysis of cells treated with pimasertib at cytotoxic (5 µM, A549 and Calu-6) and cytostatic (non-cytotoxic) (0.5 µM in A549) doses. RNA was obtained for analysis from two independent experiments and FKPM values for each gene were averaged prior to further analysis. (B) GO analysis for the 555 significantly altered death-associated genes. (C) Inhibitors (bold text) block the function of BCL-2-family proteins to induce apoptosis. (D) Cell death over time in A549^N^ cells co-treated with MEKis and the BCL-xL inhibitors ABT-737 or A-1155463 (both 5 µM). (E) Cell death kinetic parameters for treatments in D. (F) Cell death over time in A549^N^ cells co-treated with MEKis or bortezomib (Btz) and the MCL1 inhibitor S63845 (5 µM). (G) Cell death kinetic parameters for treatments in F. (H) Mean single cell fluorescence intensity (M.F.I.) for MCL1 and BCL-xL in individual A549^N^ cells determined by immunofluorescence at 48 h ± Pim (5 µM). Coefficient of variation (CV) is indicated. (I) MCL1 M.F.I. over time in A549^N^ cells treated as indicated. Coefficient of variation (CV) is shown for all conditions, and ranges (min-max) are shown for the 96 h sample. For H and I, at least 115 individual cells are quantified per condition from multiple independent microscopic fields, and median and interquartile ranges are indicated. (J) Expression of MCL1 in A549^N^ cells transduced with CMV-Empty and CMV-MCL1 lentivirus. (K) Cell death over time ± MCL1 overexpression, as in J. (L) D_R_ values computed from lethal fraction curves in K for MEKis. Results are from three independent experiments and represent the mean ± SD (D,F,K) or the mean ± 95% C.I. (E, G, L).

Based on literature precedent, we first examined the roles of the pro-apoptotic BH3-only protein BIM and the antiapoptotic protein BCL-xL (Cragg et al., 2008; Hata et al., 2014; Luciano et al., 2003; Meng et al., 2010). As expected, BIM levels increased, and BCL-xL levels decreased, following MEKi treatment in both A549^N^ and Calu-6^N^ cells (Figure S4C). We hypothesized that if slow FK was due to heterogenous expression of either protein, BIM overexpression or BCL-xL inhibition alone should promote the synchronization of cell death within the population. Specifically, such an effect would be apparent in higher D_R_ values in response to MEKi treatment. Doxycycline (Dox)-inducible overexpression of wild-type BIM, but not inactive BIM^G156E^ (Marani et al., 2002), did shorten the time to cell death onset (i.e. D_O_) but did not alter D_R_ in response to MEKis, across a range of tested Dox doses (Figure S4D-H). Likewise, co-treatment of A549^N^ cells with the BCL-xL/BCL-2 inhibitor ABT-737 or the BCL-xL-selective inhibitor A-1155463 shortened D_O_ without increasing D_R_ (Figure 4C-E). Thus, BIM and BCL-xL helped set the threshold for MEKi-induced cell death but were not primary regulators of the heterogeneous nature of this process between cells.

In addition to BCL-xL, the antiapoptotic protein MCL1 can also regulate apoptosis in response to inhibition of the MAPK pathway (Montero et al., 2019; Nangia et al., 2018; Pétigny-Lechartier et al., 2017; Sale et al., 2019). Interestingly, cotreatment with the selective MCL1 inhibitor S63845 shortened D_O_ but also more than doubled MEKi-induced D_R_, from less than 2% h^-1^ to over 4% h^-1^ in A549^N^ cells (Figure 4F,G). This effect was specific to MEKis, as MCL1 inhibition did not alter D_R_ in response to the proteasome inhibitor bortezomib (Figure 4F,G). Similar but quantitatively weaker effects on D_R_ were observed in S63845-treated Calu-6^N^ cells in response to MEKi treatment (Figure S4I,J). Consistent with the small molecule inhibitor data, polyclonal CRISPR/Cas9-mediated *MCL1* disruption enhanced MEKi-induced FK (Figure S4K,L, and see below). Thus, MCL1 was an important negative regulator of MEKi-induced slow FK.

Given the above results, we examined how MCL1 expression varied between individual cells, as determined using protein immunofluorescence, in response to MEKi treatment. As a comparison, we also examined the expression of BCL-xL. Consistent with our bulk analysis, Pim treatment (5 µM, 48 h) reduced single cell BCL-xL protein levels (Mann Whitney test, *P* < 0.0001) (Figure 4H). BCL-xL expression was also more homogeneous between cells following Pim treatment, as inferred from the coefficient of variation in protein expression (Figure 4H). By contrast, the levels of MCL1 increased following Pim treatment, and variation in protein expression between cells remained high (Figure 4H). Indeed, when examined in response to MEKi treatment over time, MCL1 levels became increasingly heterogeneous, as assessed either by coefficients of variation or the range of absolute MCL1 expression between remaining live cells (Figure 4I). Higher expression of MCL1 in an individual cell could inhibit cell death by allowing for greater sequestration of pro-apoptotic BH3-only proteins. Consistent with this model, enforced MCL1 overexpression significantly reduced the population death rate (i.e. D_R_) in response to MEKis (Figure 4J-L). We conclude that heterogeneous MCL1 expression between cells is an important contributor to FK in response to MAPK pathway inhibitors.

### The regulation of FK is lethal stimulus-specific

Whether the molecular mechanisms governing FK vary between lethal stimuli, or are shared, is poorly understood. Our evidence suggested that the mechanisms governing FK in response to MEKis were unlikely to be shared with other lethal molecules. Further support for this model was obtained from the analysis of a series of fifteen (sub-) clonal cell lines isolated from our starting population of A549^N^ cells, each exposed to Pim (5 µM), Tram (0.5 µM), bortezomib (Btz, 50 nM) or erastin2 (Era2, 2 µM), and examined in parallel using time-lapse imaging (Figure 5A). Substantial variability was observed been these clonal populations in MEKi-induced FK, while more consistent responses were observed in response to both Btz and Era2 (Figure 5B). This directly demonstrates that the molecular mechanisms governing FK in response to MEKis are distinct from those contributing to FK in response to other lethal stimuli. This experiment also further highlighted the utility of highly parallelized analysis of FK for comparative studies.

**Figure 5.**
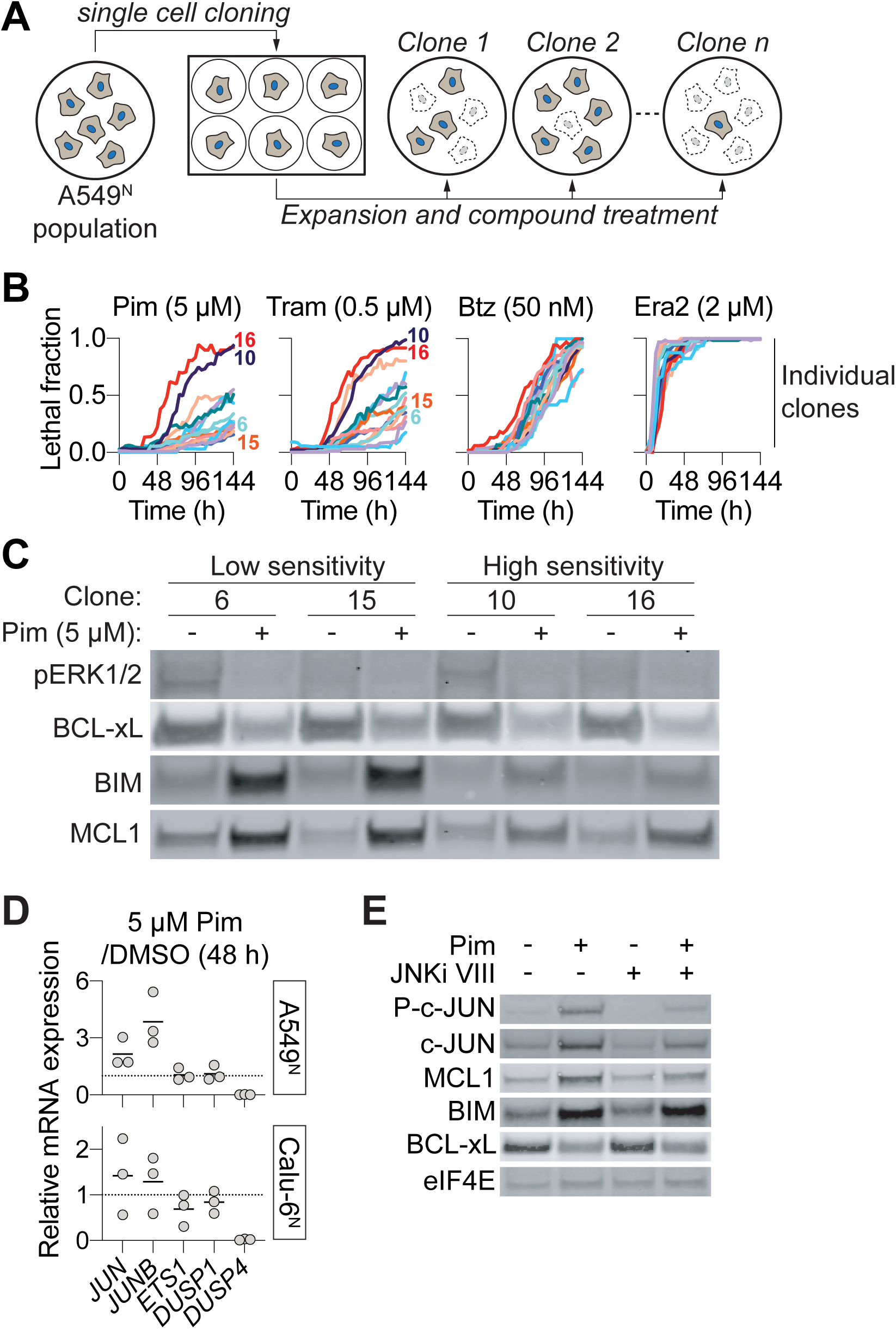
Heterogeneous MCL1 expression correlates with death sensitivity. (A) Overview of the strategy used to isolate clonal populations of A549^N^ cells. (B) Cell death over time for individual clonal populations examined in parallel. Each independent clone was derived from a single cell, expanded, and tested a single time. Key clones analyzed at the protein level in C are numbered. (C) Expression of key apoptotic proteins in low and high sensitivity clones. (D) RT-qPCR analysis in A549^N^ cells of selected IEGs. Individual datapoints represent individual experiments, and mean values are indicated by the horizontal bar. (E) Western blot of A549^N^ cells. Pim and JNKi VIII were both used at 5 µM.

These clonal cell lines also provided an opportunity to study how differences in MEKi-induced FK related to the expression of MCL1. Based on our existing data (Figure 4H-L), we hypothesized that high levels of MCL1 expression would correlate with relative resistance to MEKi-induced FK. To test this hypothesis, we picked clones that exhibited relatively low (#6 and #15) or high (#10 and #16) sensitivity to MEKi-induced FK and examined protein expression. Both low and high sensitivity clones exhibited similar basal phosphorylated ERK1/2 and BCL-xL expression, and similar loss of both species following MEKi treatment (Figure 5C). By contrast, low sensitivity clones accumulated substantially higher levels of MCL1 and BIM following MEKi treatment compared to high sensitivity clones (Figure 5C). Thus, high levels of MCL1 accumulation correlates with slower MEKi-induced FK, possibly due to a greater capacity to ‘buffer’ the accumulation of pro-death proteins like BIM. The ability to track FK over time in multiple independent populations was central to elucidating this relationship.

The pathway linking MAPK pathway inhibition to increased MCL1 levels was unclear. In our RNA-seq analysis we observed increased expression of several JUN-family immediate early genes, consistent with activation of the JNK pathway (Figure S5). This was intriguing as in other systems JNK1/2 can suppress apoptosis by phosphorylating and stabilizing MCL1 (Hirata et al., 2013; Kodama et al., 2009). We confirmed activation of the JNK pathway in response to MEKi treatment by RT-qPCR and Western blotting for JUN family members in parental A549^N^ cells (Figure 5D,E). Moreover, we observed that inhibition of JNK function using the small molecule inhibitor JNKi VIII (Miura et al., 2018) inhibited JUN expression and phosphorylation, and specifically blocked MCL1 accumulation in response to MEKi treatment, without affecting BCL-xL or BIM levels (Figure 5E). Thus, inhibition of the MAPK pathway can cause of activation of the stress responsive JNK pathway and stabilization of anti-apoptotic MCL1 protein, impacting the kinetics of FK.

### Compound interactions alter MEKi-induced fractional killing over time

In the clinic, few drugs are employed as single agents and it is therefore of interest to understand the impact of drug interactions specifically on FK (Palmer et al., 2019). We show, above, that it is possible to detect chemical interactions that enhance FK (e.g. MEKis + MCL1 inhibition, Figure 4F). Our ability to monitor FK in many populations in parallel suggested provided a means to search for modulators of FK more systematically. As proof-of-principle, we examined the effect on Pim-induced cell death of 261 structurally and functionally distinct compounds in A549^N^ cells. The ability of all 261 compounds to modulate Pim-induced cell death was computed using an analytic framework based on the Bliss model of drug interactions (Conlon et al., 2019). This dataset provided an opportunity to systematically assess how drug interactions impacted FK (Figure 6A). Strikingly, multiple ATP-competitive and allosteric mechanistic target of rapamycin (mTOR) inhibitors potently accelerated Pim-induced FK, both reducing D_O_ and increasing D_R_ by as much as five-fold (Figure 6B). In follow-up experiments we confirmed that two structurally and mechanistically distinct mTOR inhibitors, the ATP competitive inhibitor AZD8055 and the rapalog rapamycin, both accelerated MEKi-induced FK specifically, with no sensitization observed towards bortezomib (Figure 6C). These results indicate that it is possible to pinpoint drug interactions that specifically alter FK in a high-throughput manner.

**Figure 6.**
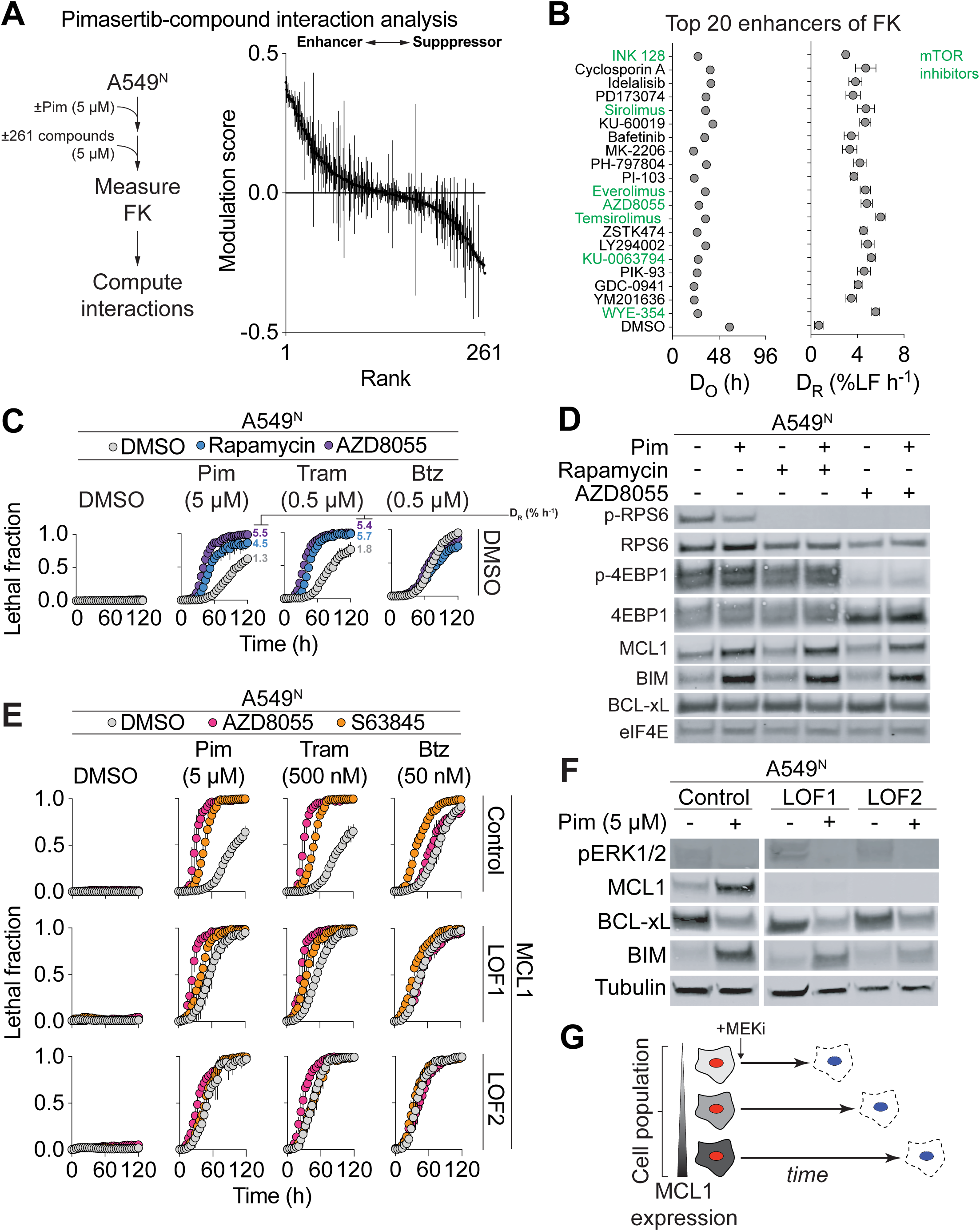
High-throughput analysis of drug interactions impacting fractional killing. (A) A drug-drug interaction screen in Pim-treated A549^N^ cells. Cells were co-treated ± 5 μM Pim plus a library of 261 bioactive compounds and controls. Data on the y-axis represent deviations of the observed normalized area under the curve (nAUC) value from the expected nAUC value for each library compound in combination with Pim as calculated using the Bliss model of drug independence (see Methods). The screen was performed three separate times and the results represent mean ± SD. (B) Kinetics of cell death computed for the top 20 enhancers of Pim-induced cell death. mTOR inhibitors are indicated in green. (C) Cell death over time for the indicated combinations. Rapamycin was used at 50 nM, and AZD8055 at 500 nM. Mean D_R_ values were computed from lethal fraction curves and are indicated. (D) Analysis of protein expression by Western blotting ± Pim (5 µM), rapamycin (50 nM) or AZD8055 (500 nM) at 48 h. (E) Cell death over time in Control and two independent MCL1 loss of function clones (LOF1/2). (F) Protein expression in Control, and MCL1 LOF1 and LOF2 cell lines. Compound treatment was for 48 h. (G) Model expressing how variability in MCL1 expression links to variable membrane permeabilization (i.e. cell death) over time. Data are from three independent experiments and represent mean ± 95% C.I. (B) or mean ± SD (C,E).

Our finding that mTOR inhibition accelerated MEKi-induced FK was intriguing in light of evidence that mTOR signaling promotes *MCL1* mRNA expression and post-translational MCL1 stability (Koo et al., 2015; Mills et al., 2008). Based on these findings, we hypothesized that mTOR inhibition accelerated MEKi-induced FK through by reducing MCL1 expression. However, mTOR inhibitors did not reduce basal MCL1 expression or prevent Pim treatment from increasing MCL1 protein levels in A549^N^ cells (Figure 6D). We confirmed that mTOR inhibitors blocked the phosphorylation of the canonical downstream mTOR targets RPS6 (both) and/or 4EBP1 (AZD8055 only) and did not alter the effect of Pim on BIM upregulation or BCL-xL downregulation (Figure 6D).

These results suggested that mTOR signaling might promote FK independent of effects on MCL1. To investigate further, we generated two clonal *MCL1* gene disrupted cell lines, one (LOF1) that appeared to be a severe loss of function, and one (LOF2) that appeared to be a complete knockout, based on MCL1 protein expression and sensitivity to the MCL1 inhibitor S63845 (Figure 6E,F). Consistent with results obtained using the polyclonal CRISPR method (Figure S4K,L), MEKi-induced FK was substantially accelerated by clonal disruption of MCL1 (Figure 6E). Notably, mTOR inhibition enhanced MEKi-induced FK equivalently in the presence or absence of MCL1. This suggests that mTOR inhibition likely sensitizes to MEKi-induced FK independent of MCL1. These results demonstrate that it is possible to systematically search for and isolate compound-compound interactions that specifically alter the kinetics of FK.

## DISCUSSION

Heterogeneous cellular phenotypes within one population are common in nature, with drug-induced fractional killing in tumors being one clinically important manifestation (Bigger, 1944; Inde and Dixon, 2018; Niepel et al., 2009; Palmer et al., 2019). Quantifying population-level cell heterogeneity can be laborious, especially when tracking the fate of hundreds of individual cells within a single population. We find that population-level time-lapse measurements of live and dead cells enables FK to be effectively quantified without tracing individual cells. This enables the quantitative, large-scale analysis of FK in hundreds of populations in parallel in a single experiment. A limitation of this approach is that high-throughput time-lapse imaging remains less accessible than more traditional methods. However, we show that traditional bulk biochemical methods (e.g. CellTiter-Glo and AlamarBlue) suffer limitations that make them unsuitable for the precise quantification of FK between conditions.

Consistent with previous single-cell analyses of different individual populations (Miura et al., 2018; Paek et al., 2016; Roux et al., 2015; Spencer et al., 2009) we find that FK is fundamentally a kinetic phenomenon. At any given time, some fraction of cells within the population may be alive and others dead. Our findings make clear that at any given timepoint this fraction can vary substantially between lethal treatments and genetic backgrounds. It is important to note that for many lethal conditions all cells within the population would ultimately die, at some timepoint. What these methods enable is for the heterogeneity of cell death initiation between cells in population to be easily quantified and compared. In turn, this allows for the contribution of different molecular mechanisms to the kinetics of cell death to be specifically isolated. This contrasts with the majority of mechanistic studies, which seek only to arrive at a binary (yes/no) distinction concerning whether a given protein or mechanism contributes to a lethal process at any timepoint.

As a case study to search for molecular regulators of FK, we focused on inhibitors of the MAPK pathway, finding that MCL1 plays an important role in this context. MCL1 expression is inherently highly variable, likely due to the short half-life of *MCL1* mRNA and the instability of the protein (Senichkin et al., 2020). Our pharmacological, imaging, and overexpression studies suggest that differences in the initiation of between cells in the population are linked to variation between cells in MCL1 protein expression, with the ability to accumulate higher levels of MCL1 associated with later onset of cell death within the population (Figure 6G). At the molecular level, this may relate to differences in the time it takes for the entire MCL1 pool within a given cell to be fully overcome by the expression of pro-death BH3 proteins like BIM. Notably, and unlike inhibition of BCL-xL, pharmacological or genetic inactivation of MCL1 ‘homogenizes’ the response of individual cells to MAPK pathway inhibition, resulting in an effective acceleration of FK. That said, MCL1 inactivation does not render cells uniformly sensitive to death at the exact same time (i.e. D_R_ remains under 10% h^-1^). One goal of future work is to define MCL1-independent mechanisms governing FK in response to MAPK pathway inhibition. Evidence from our compound interaction study suggests that one such mechanism involves mTOR, perhaps through regulation of a pro-apoptotic protein like BAD (Harada et al., 2001).

We find that the kinetics of FK differ substantially between drugs in the same genetic background and for the same drug in different backgrounds. Two lines of evidence suggest a single molecular mechanism is unlikely to account for these differences. First, proteins or mechanisms that regulate FK-like phenomena in other contexts, including p53, metabolism, and the cell cycle (Lagadinou et al., 2013; Miura et al., 2018; Paek et al., 2016; Roux et al., 2015), did not modulate FK in response to MEKis. Second, a single A549 population contains diverse sub-populations of cells with similar sensitivity to proteasome inhibition or nonapoptotic cell death induced by erastin2, but distinct sensitivities to MEKis. Thus, FK cannot be governed by a single unified mechanism. Further investigation of FK using the large-scale comparative approach presented here could help elucidate the constellation of different molecular mechanisms that regulate FK and provide new insights into this important form of phenotypic heterogeneity.

## STAR♦METHODS

Detailed methods are provided in the online version of this paper and include the following:

- KEY RESOURCES TABLE
- LEAD CONTACT AND MATERIALS AVAILABILITY
- EXPERIMENTAL MODEL AND SUBJECT DETAILS
  - Cell lines and Culture Conditions
  - Chemicals and Reagents

- METHOD DETAILS
  - Measurement of cell viability using CellTiter-Glo and PrestoBlue
  - Lentivirus generation and infection
  - Generation of CRISPR/Cas9 knockout cell lines
  - Isolation of clonal cell populations
  - Cell seeding and compound treatment
  - Doxycycline-inducible protein overexpression
  - CMV vector protein overexpression and rescue
  - Time-lapse imaging, LED curve fitting and parameter extraction
  - Kinase translocation reporter (KTR) analysis
  - Cell metabolism and ROS analysis
  - Western blotting
  - RNA-seq
  - Propidium iodide cell cycle profiling
  - Immunofluorescence imaging
  - Chemical library screening

- QUANTIFICATION AND STATISTICAL ANALYSIS
- DATA AND CODE AVAILABILITY

## SUPPLEMENTAL INFORMATION

Supplemental information includes four figures and can be found with this article online.

## SUPPLEMENTAL INFORMATION

Supplemental information includes four figures and can be found with this article online.

## ACKNOWLEDGEMENTS

We thank L. Magtanong for comments, A. Gitler for plasmids and B. Stockwell for erastin. S.J.D. is supported by the NIH (1R01GM122923) and a Damon Runyon-Rachleff Innovation Award.

## AUTHOR CONTRIBUTIONS

Conceptualization, Z.I., G.C.F., and S.J.D.; Methodology, Z.I., G.C.F.; Investigation, Z.I., G.C.F., and K.D.; Writing – Original Draft, Z.I. and S.J.D.; Writing – Review & Editing, Z.I., G.C.F., and S.J.D.; Funding Acquisition and Supervision, S.J.D.

## STAR♦METHODS

### Lead Contact for Materials Availability

Further information and requests for resources and reagents should be directed to and will be fulfilled by the Lead Contact, Scott Dixon.

### Experimental Model and Subject Details

#### Cell lines and culture conditions

The following cell lines were obtained from ATCC (Manassas, VA): A549 (CCL-185, sex: male), Calu-6 (HTB-56, sex: female), NCI-H1299 (CRL-5803, sex: male), NCI-H2291 (CRL-5939, sex: male), T98G (CRL-1690, sex: male), U-2 OS (HTB-96, sex: female) and HEK 293T cells (CRL-3216, sex: female). Cell lines stably-expressing nuclear-localized mKate2 (denoted by the superscript ‘N’) were generated by infection with lentivirus (Cat# 4625, Essen BioScience, Ann Arbor, MI) followed by puromycin (10 µg/mL, Cat# A11138-03, Life Technologies, Carlsbad, CA) selection for three days. For kinase translocation reporter (KTR) and polyclonal CRISPR cell lines described below, mKate2-expressing cell lines were generated with a bleomycin-selectable lentiviral construct (Cat# 4627, Essen BioScience) to allow subsequent infection with puromycin-selectable KTR constructs. After infection with bleomycin-selectable lentivirus, cell lines were sorted to isolate mKate2-positive cells using a FACSAria II Fluorescence Activated Cell Sorter (BD Biosciences, San Jose, CA) at the Stanford Shared FACS Facility. Cell lines were cultured in an appropriate base media supplemented with 10% fetal bovine serum (Cat# 26140-079, Thermo Fisher Scientific) and 1% penicillin/streptomycin (Cat# 105070-063, Life Technologies). Base media used was DMEM (Cat# MT-10-013-CV, Thermo Fisher Scientific) for A549, Calu-6, NCI-H1299 (H1299), T98G, and HEK293T, RPMI (Cat# SH30027FS, Thermo Fisher Scientific) for NCI-H2291, and McCoy’s 5A (Cat# MT-10-050-CV, Thermo Fisher Scientific) for U-2 OS. Cell lines were cultured in incubators at 37°C and 5% CO_2_. For passaging, cells were rinsed with HBSS (Cat# 14025-134, Life Technologies), dissociated with 0.25% trypsin-EDTA (Cat# SH3004201, Thermo Fisher Scientific), quenched with the corresponding complete media, and counted using a Cellometer Auto T4 Bright Field Cell Counter (Nexcelom, Lawrence, MA). Upon counting, cells were subsequently seeded for experiments as described below.

#### Chemicals and reagents

Camptothecin (Cat# S1288), vinblastine (Cat# S1248), pimasertib (Cat# S1457), trametinib (Cat# S2673), ABT-737 (Cat# S1002), A-1155463 (Cat# S7800), Nutlin-3 (Cat# S1061) were purchased from Selleck Chemicals (Houston, TX). Bortezomib (Cat# NC0587961), etoposide (Cat# ICN19391825), MG-132 (Cat# 17-485), and Q-VD-OPh (Cat# OPH00101M) were purchased from Thermo Fisher Scientific. N-acetylcysteine (Cat# A8199), thapsigargin (Cat# T9033), tunicamycin (Cat# T7765), paclitaxel (Cat# T7191), JNK Inhibitor VIII (Cat# 420135), 2-deoxyglucose (Cat# D8375), oligomycin (Cat# O4876), and cycloheximide (Cat# C7698) were obtained from Sigma-Aldrich (St. Louis, MO). S63845 (Cat#21131) was obtained from Cayman Chemical (Ann Arbor, MI). Staurosporine (Cat# A8192) was obtained from ApexBio (Houston, TX). Erastin was the kind gift of Brent Stockwell (Columbia University). Erastin2 (compound 35MEW28 in (Dixon et al., 2014)) and ML162 (CAS: 1035072-16-2) were synthesized by Acme Bioscience (Palo Alto, CA). Chemical screening was conducted as described below; the library of 261 bioactive compounds was obtained from Selleck Chemicals (Cat# L2000).

## Method Details

### Measurement of cell counts and cell viability using CellTiter-Glo and PrestoBlue

U-2 OS^N^ and T98G^N^ cells were seeded into parallel 384-well plates at a concentration of 1,500 cells/well, briefly spun at 500 RPM to settle cells to the bottom of plates, and then incubated for 24 h in a tissue culture incubator prior to the start of the experiment. The next day, 4 μL of 10X drug stock prepared in media was added to 36 μL of fresh media to a final concentration of 1X. At 72 h, both plates were scanned for mKate2 positive objects (i.e. live cells) using an IncuCyte imaging system (Essen BioScience) and the imaging parameters described in Table S1. Then immediately after the scans, one plate was subjected to CellTiter-Glo (CTG, Promega, Cat# G7570) analysis and the other to Presto Blue (PB, Thermo Fisher Scientific, Cat# A13262) analysis using the above commercially available reagents. For CTG analysis, cells were lysed by addition of 40 μL of CellTiter-Glo reagent for two min while shaking. Cells were allowed to equilibrate for 10 min post-lysis followed by luminescence reading using a Cytation3 multimode plate reader (BioTek, Winooski, VT, USA) set to luminescence, optics position = top, gain = 135. For PB analysis, 10 μL of 5X PB reagent was added to each well and the plate was then allowed to incubate at 37°C and 5% CO_2_ for 30 min, followed by fluorescence reading using a Cytation3 multimode plate reader (BioTek, Winooski, VT, USA) set to ex/em 560/590, optics position = bottom, gain = 75.

### Lentivirus generation and infection

For generation of lentivirus, HEK293T cells were seeded at 300,000 cells per well in six well plates (Cat# 07-200-83, Thermo Fisher Scientific) one day before transfection. Plasmids for lentiviral transduction were transfected into HEK293T cells along with lentiviral packaging plasmids pCMV-VSV-G, pMDLg/pRRE, and pRSV-Rev (obtained from Dr. Michael Bassik, available via Addgene as Cat# 8454, 12251, and 12253). Transfection was conducted using PolyJet Transfection Reagent (Cat# SL100688, SignaGen Laboratories, Rockville, MD) according to manufacturer instructions, using a plasmid mix of .25 μg pCMV-VSV-G, 0.25 μg pMDLg/pRRE, 0.25 μg pRSV-Rev, and 0.75 μg of the appropriate transfer plasmid. Lentivirus-containing supernatant from the transfected cells was harvested 48 and 72 hours after transfection and filtered through a 0.45 μm Millex filter (Cat# SLHV033RS, EMD Millipore, Burlington, MA) using a 10 mL syringe (Cat# 14-817-30, Thermo Fisher Scientific). Filtered lentivirus was stored at −80°C or used immediately. For infection, cell lines were seeded a day before infection in 12 well plates (Cat# 07-200-82, Thermo Fisher Scientific) to be approximately 50% confluent on the day of infection. On the day of infection, supernatant was removed from the cells and replaced with 500 μL of lentivirus-containing media and 500 μL of polybrene (Cat# H9268-5G, Sigma-Aldrich, St. Louis, MO) diluted to a final concentration of 16 μg/mL in the appropriate complete media. Selection with the appropriate antibiotic was begun after two days.

### Generation of CRISPR/Cas9 knockout cell lines

Two methods were used to generate CRISPR/Cas9 knockout cell lines, one yielding polyclonal knockout lines and the other clonal knockout cell lines. Polyclonal knockout cell lines were generated by lentiviral infection of cell lines with the lentiCas9-Blast plasmid and an MCL1 sgRNA-expressing plasmid. lentiCas9-Blast was a gift from Feng Zhang (Addgene plasmid #52962; RRID: Addgene_52962). sgRNA expressing plasmids were produced by cloning sgRNA oligonucleotides with the appropriate adaptor sequences (see Table S2) into plasmid pMCB306 after digesting the plasmid with restriction enzymes BstXI and Bpu1120I (Cat# FD1024, FD0094, Thermo Fisher Scientific). pMCB306 was a gift from Michael Bassik (Addgene plasmid #89360; RRID: Addgene_89360). After infection and antibiotic selection (10 μg/mL blasticidin or puromycin for three days), knockout cell lines were validated by Western blot.

Clonal CRISPR/Cas9 knockout cell lines were generated as described previously (Cao et al., 2019). Briefly, sgRNA expressing plasmids were produced by cloning sgRNA oligonucleotides with the appropriate adaptor sequences (see Table S2) into plasmid spCas9-2A-EGFP (PX458). spCas9-2A-EGFP (PX458) was a gift from Feng Zhang (Addgene plasmid #48138; RRID: Addgene_48138). Knockout cell lines were generated from mKate2-expressing A549 cells generated with a bleomycin-selectable lentiviral construct, allowing for subsequent rescue using a puromycin-selectable construct. Cells were seeded and transfected with the sgMCL1-expressing plasmid in a 6 well plate using Lipofectamine LTX transfection reagent (Cat# 15338030, Thermo Fisher Scientific) according to manufacturer’s instructions. 2 days after transfection, single cells were isolated and sorted into a 96 well plate using an Influx Special Order fluorescence activated cell sorter (BD Biosciences, San Jose, CA) at the Stanford Shared FACS Facility. Cells were sorted into DMEM media supplemented with 30% fetal bovine serum (Cat# 26140-079, Thermo Fisher Scientific) and 1% penicillin/streptomycin (Cat# 105070-063, Life Technologies). Clones were monitored and expanded to larger formats over the following 4 weeks, then validated for MCL1 knockout by Western blot.

### Isolation of clonal cell populations

Single-cell clones were isolated and expanded from mKate2-expressing A549 cells. Low passage cells were trypsinized and brought to the Stanford Shared FACS facility for single cell sorting as described for CRISPR clone sorting above. Two sets of twelve clones were passaged into new 96 well plates on days 23 and 25 after sorting. Each set of clones was treated with lethal drugs the day after seeding and monitored by Incucyte for 144 hours. Clones with insufficient cell number or heterogeneous expression of mKate2 were excluded from analysis. When seeded for drug treatment, clones were also expanded to obtain cell lysates for protein expression analysis. Cells were harvested and lysed for Western Blotting as described below, and protein expression was compared across clones.

### Cell seeding and compound treatment

Experiments measuring death after treatment with MEK inhibitors (or other lethal compounds), either alone or in combination with modulators (e.g. CHX, Q-VD-OPh, or Bcl-2 family inhibitors), were conducted as follows. Cells were seeded one day prior to drug addition in tissue culture treated clear-bottom 96- or 384-well plates (Fisher Scientific Cat# 07-200-588, 07-201-013). Appropriate cell seeding density was determined for each cell line to ensure seeding dense enough for cells to proliferate normally and sparse enough to minimize death due to overcrowding before the end of the experiment. On the day of treatment, drugs were diluted to final doses in the appropriate cell culture media noted above. Seeding media was removed from the plate and replaced with drug-containing media. For experiments to be analyzed by STACK, 20 nM SYTOX Green dye (Life Technologies, Cat# S7020) was included in the treatment media, and drug treated plates of cells were imaged every 4 h using an IncuCyte. Images were analyzed automatically as described below. For drug cycling experiments, cells were seeded in 384-well plates. On the day of drug removal, a multichannel pipette was used to remove the drug-containing media and wash once with normal media before adding normal media to the wells for the drug-untreated period.

### Doxycycline-inducible protein overexpression

To generate doxycycline-inducible cell lines, cells were infected with lentivirus carrying the pLenti CMV rtTA3 Blast (w756-1) plasmid and selected with blasticidin (Cat# A1113902, Thermo Fisher Scientific). Genes of interest were cloned into the pLenti CMVTRE3G eGFP Neo (w821-1) plasmid using a Gibson Assembly Master Mix kit (Cat# E2611S, New England Biolabs, Ipswich, MA). pLenti CMV rtTA3 Blast (w756-1) and pLenti CMVTRE3G eGFP Neo (w821-1) were gifts from Eric Campeau (Addgene plasmids #26429 and #27569; RRIDs: Addgene_26429 and Addgene_27569). *BCL2L1* (Bcl-xL) and both wild type and G156E-mutant *BCL2L11* (BIM) open reading frames for Gibson Assembly were obtained as custom synthesized gBlocks from Integrated DNA Technologies (Coralville, IA). Lentivirus was generated as described above and rtTA-expressing cell lines were infected and subsequently selected with 10 μg/mL Geneticin (Thermo Fisher Scientific, Cat# 11811-031); complete selection with Geneticin required three passages in antibiotic. An uninfected control population was treated with Geneticin and passaged in parallel to ensure complete selection.

### CMV vector protein overexpression and rescue

CMV overexpression of MCL1 and rescue of MCL1^KO^ cells was carried out using the pLenti CMV Puro DEST backbone. pLenti CMV Puro DEST (w118-1) was a gift from Eric Campeau & Paul Kaufman (Addgene plasmid# 17452; RRID: Addgene_17452). The open reading frame for *MCL1* was obtained as a gift from Dr. Aaron Gitler as a pDONR plasmid from the human ORFeome. The MCL1 ORF was cloned into the pLenti CMV vector using a Gateway LR Clonase II kit (Cat# 11791-020, Thermo Fisher Scientfic, Waltham, MA). The resulting plasmid is missing a stop codon; site-directed mutagenesis was carried out using the Q5 Site-Directed Mutagenesis Kit (New England Biolabs, Cat# E0554S) to introduce a stop codon. Further site-directed mutagenesis was conducted to introduce phosphosite mutations for comparative rescue experiments in MCL1 KO cells. Mutagenesis primers are listed in Table S2. Lentivirus was generated as described above, generating virus with both the CMV-MCL1 wild type and mutant plasmids and the control backbone plasmid that did not undergo Gateway recombination. Cell lines were infected with the harvested virus, and infected cells were selected with 5 μg/mL puromycin for three days.

### Time-lapse imaging, LED curve fitting and parameter extraction

Image analysis was conducted using Incucyte Zoom software (Essen BioScience). Image analysis parameters were optimized for each cell line; parameters are listed in Table S1. Lethal fraction scoring, LED curve fitting and parameter extraction were carried out as described previously (Forcina et al., 2017). Red and green object counts (as quantified according to the parameters in Table S1) were used to the compute the lethal fraction at each timepoint and then using Prism 7 software (GraphPad, San Diego, CA), lethal Fraction curves were fit using the “Plateau followed by one-phase association” function to obtain an LED fit. D_O_ (corresponding to X0 in Prism) and D_R_ (corresponding to K in Prism) values were extracted from the successful curve fits. For parameter analysis in Figure 2, only lethal compound conditions yielding average lethal fraction scores > 0.5 across all three independent experiments were examined.

### Kinase translocation reporter (KTR) analysis

Kinase translocation reporter (KTR) cell lines were generated by lentiviral transduction of the pLentiCMV Puro DEST ERKKTRClover and pLentiPGK Puro DEST p38KTRClover plasmids (gift of Dr. Markus Covert, obtained via Addgene, Cat# 59150, 59152). Infected cells were selected with 10 μg/mL puromycin for 3 days. For drug treatment experiments, periodic IncuCyte imaging started the day after seeding and continued for 24 hours before drug addition to establish an untreated baseline. No more than 30 min before the 24 h timepoint, seeding media was removed from the cells and drug-containing media was added to the plate. To analyze the collected images, Zoom analysis parameters were modified as listed in Table S1. KTR reporter was detected in the green channel in lieu of SYTOX Green in these experiments. mKate2 signal marked cell nuclei, and cells in which the KTR reporter translocated to the nucleus produced a dual-positive overlap signal that was quantified in the software.

### Cell metabolism and ROS analysis

For analysis of glycolytic and oxidative metabolism upon drug treatment, A549N cells were seeded in normal media in Seahorse xFP plates (Cat# 103022-100, Agilent Technologies, Santa Clara, CA) 24 h prior to conducting the assay. Sensor cartridges were also hydrated on the preceding day. On the day of the assay, cells were washed with complete Seahorse XF media (Cat# 103334-100, Agilent Technologies) supplemented to final concentrations of 25 mM glucose, 1 mM pyruvate, and 4 mM glutamine and adjusted to pH 7.4. After washing twice and replacing media with complete XF media, plates were imaged by IncuCyte to obtain cell counts for normalization and the plate was incubated in a non-CO2 incubator for one hour to de-gas. 10X solutions of drugs for injection (oligomycin, 2-DG, and ethanol vehicle control) were loaded into the sensor cartridge, and the cell plate and sensor cartridge were inserted into a Seahorse xFP Analyzer (Agilent Technologies) to obtain extracellular acidification rate (ECAR) and oxygen consumption rate (OCR) metrics. ECAR and OCR values were measured automatically approximately every 6 minutes; after four baseline measurements, drugs were injected from the cartridge and metabolic parameters were measured continuously over the remainder of the experiment.

For FACS analysis of ROS levels, cells were trypsinized and resuspended in HBSS containing 20 uM H_2_DCFDA (Thermo Fisher Scientific, Cat# D-399). Cells were incubated in a 37°C, 5% CO_2_ incubator for 30 min to stain, then spun down at 1000 rpm for 5 min. H_2_DCFDA containing supernatant was aspirated, and the pellet of stained cells was resuspended in HBSS and strained through a 35 μm mesh filter into a test tube (Corning, Cat# 352235) for sorting. Cells were sorted on a FACSAria II Fluorescence Activated Cell Sorter (BD Biosciences, San Jose, CA) at the Stanford Shared FACS Facility. The top and bottom 5% of cells based on H_2_DCFDA were sorted into DMEM, counted, and seeded in 384 well plates prepared with diluted drugs before the sort. IncuCyte imaging began the following day and images were acquired once every 24 h.

### Western blotting

Cells for Western blot analysis were treated in 6 well plates (Thermo Fisher Scientific, Cat# 07-200-83) or 10 cm dishes (USA Scientific, Ocala, FL, Cat# CC7682-3394). At the time of harvest, treatment media was removed, and cells were washed with ice cold HBSS and detached with cell lifters (Fisher Scientific, Cat# 07-200-364). Detached cells in HBSS were spun down at 1000 rpm for 5 min, then supernatant was removed, and the remaining pellets were frozen overnight at −80°C. NP-40 buffer (1% NP-40 detergent, 50 mM Tris-Cl pH 8.0, 150 mM NaCl) was used for lysis. NP-40 buffer was supplemented with protease/phosphatase inhibitor cocktail (Cell Signaling Technology, Danvers, MA, Cat# 5872S) and used to lyse frozen pellets. Lysis proceeded for 45 min on ice, then samples were sonicated for 10 pulses of 1 second each at 50% amplitude with a Fisherbrand Sonic Dismembrator (Fisher Scientific, Cat# FB120110). Lysates were centrifuged for 15 min at 12,700 rpm to remove debris, then protein concentration was measured by BCA assay kit (Thermo Fisher Scientific, Cat# 23252).

Lysates were combined with Bolt sample buffer and reducing agent (Thermo Fisher Scientific, Cat# B0007, B0009) and heated for 10 min at 70°C. Prepared samples were loaded onto Bolt 4-12% gradient gels (Thermo Fisher Scientific, Cat# BG04120BOX) and run for 1 hour 45 min at 100 mV. Protein from the finished gel was transferred to a nitrocellulose membrane using an iBlot 2 Transfer stack (Invitrogen, Cat# IB23002). Membranes were washed in Odyssey Blocking buffer (LI-COR Biosciences, Lincoln, NE) at room temperature for 1 hour, then incubated in primary antibody mixture overnight at 4°C. Primary antibody incubation was followed by three, seven min washes in Tris-Buffered Saline + 0.1% Tween (TBS-T), then a 45 min room temperature incubation in secondary antibody mix. Membranes were washed three more times in TBS-T before imaging on an Odyssey CLx imaging system (LiCOr Biosciences). In cases where antibodies against both the phopho and total protein were used, both forms of the protein were blotted sequentially on the same membrane. The phosphorylated version of the protein was blotted first; after imaging, the membrane was stripped with a 20 min wash in NewBlot Nitro Stripping Buffer followed by three TBS-T washes. Membranes were then reprobed by the same protocol using an antibody mix targeting the total protein.

### RNA-seq

RNA was harvested from A549 cells grown in six-well dishes. 150,000 cells were seeded into each well. The next day, cells were treated with either vehicle control (DMSO), a cytostatic, non-lethal concentration of pimasertib (0.5 µM) or a cytotoxic, lethal concentration of pimasertib (5 µM). Calu-6 cells were seeded at 300,000 cells per well and the next day treated with DMSO or Pim (5 µM). Treatments were carried out for 96 h prior to cell harvest. In vehicle-treated conditions, cells were split 1:8 after 24 hours of treatment to prevent overgrowth. Note: for lethal conditions (i.e. 5 µM Pim), treatment resulted in ∼50% cell death at the time of cell harvest. At the time of harvest, treatment media was removed, and cells were washed with ice cold HBSS (to remove debris) and detached with cell lifters. Detached cells were pelleted (1000 rpm x 5 min), the HBSS supernatant was removed, and the cell pellets frozen at −80°C overnight. The next day, cells were lysed and RNA extracted using a Qiashredder Kit and RNeasy Plus Mini Kit (Qiagen, Cat# 796554, 74134). Purity, concentration, and integrity of RNA were assessed by NanoDrop Spectrophotomer (Thermo Fisher Scientific) and by Eukaryote Total RNA Nano chip analysis on a Bioanalyzer (Agilent) at the Stanford Protein and Nucleic Acid Facility. By Bioanalyzer, the RNA Integrity Number for each sample met a threshold of at least 6.8, and NanoDrop 260/280 and 260/230 ratios met a threshold of at least 1.95. After quality control, samples were shipped on dry ice to Novogene (Sacramento, CA) for library generation and 20M read PE150 sequencing on an Illumina HiSeq 4000 platform. Bioinformatic analysis was performed by Novogene. Reads with adaptor contamination, >10% uncertain nucleotides, or >50% nucleotides with base quality <20 were filtered out, and the remaining clean reads (at least 98% of reads in all conditions) were aligned to the reference genome using Tophat v2.0.12. At least 91% of clean reads were successfully mapped in all conditions. Pearson correlations between biological replicates yielded R^2^ values above 0.98 in all conditions. Gene expression analysis was performed using DESeq 1.10.1; we isolated all gene sequences that were significantly altered (adjusted *P* value < 0.05).

### Propidium iodide cell cycle profiling

Cell cycle profiling of A549 cells was performed after treatment with drugs in a 6 well plate as described above. At the end of drug treatment, cells were washed with HBSS, trypsinized, and quenched, keeping the drug treatment media and HBSS wash for each condition to avoid loss of any loosely adherent dividing cells. Cells from each condition were spun down (500 x g, 5 min), resuspended in HBSS for one wash, then spun down again. Each pellet was resuspended in 400 µL HBSS, then 800 µL ice cold 100% ethanol was added for fixation. Cells were fixed at 4 degrees C for at least 1 hour and up to 3 weeks. When ready to be stained, fixed cells were equilibrated to room temperature, then pelleted and resuspended in HBSS containing 50 µg/mL propidium iodide (Thermo Fisher Scientific, Cat# P3566) and 500 U/mL RNAse A (Thermo Fisher Scientific, Cat# EN0531). Staining was carried out for 30 minutes at 37 degrees C, then cells were spun down, resuspended in HBSS for one wash, then spun down again. For analysis, cells were resuspended in HBSS and run on a FACSCalibur flow cytometer (BD Biosciences). Histograms of signal from the FITC channel were analyzed in FlowJo 10.6.1, using the Cell Cycle platform to estimate percentages of cells in each cell cycle phase based on propidium iodide signal.

### Immunofluorescence imaging

A549 cells were seeded in 12-well plates for immunofluorescence and treated with drugs as described above. When seeding for immunofluorescence, 12 mm round glass #1.5 coverslips (Electron Microscopy Sciences, Hatfield, PA, Cat# 72230-01) coated with poly-D-lysine (Sigma-Aldrich, Cat# P0899) were placed in each well of the 12 well plate, allowing cells to adhere to the coverslips before treatment. Once drug treatment was complete, drug-containing media was removed and cells were washed once with PBS. PBS was removed and replaced with ice cold methanol (Fisher Scientific, Cat# A935-4) to fix cells; fixation was carried out at −20°C for at least one hour and up to 72 h. To stain fixed cells, methanol was removed and replaced with PBS for one wash, then coverslips were blocked in PBS containing 3% bovine serum albumin (Gemini Bio-Products, West Sacramento, CA, Cat# 700-100P) 0.1% Triton X-100 (Sigma-Aldrich, Cat# X-100), and 0.02% sodium azide (Sigma-Aldrich, Cat# S2002) (PBS-BT). Blocking was carried out for 1 hour at room temperature or overnight at 4°C. After blocking, coverslips were transferred to a light-blocking, parafilm-coated humid chamber for staining. All staining solutions were made in PBS-BT, and all subsequent steps were carried out at room temperature. Primary antibody mixes were applied to coverslips for one hour, followed by 3 washes with PBS-BT. Secondary antibodies (Goat anti-rabbit Alexa Fluor 568 and Goat anti-mouse Alexa Fluor 488, Life Technologies, Cat# A11036 and A11029) were applied subsequently for one hour, followed by an additional 3 PBS-BT washes. Finally, nuclear staining was carried out by applying 100 ng/mL DAPI (Thermo Fisher Scientific, Cat# D1306) for 15 min, followed by 5 washes with PBS-BT. Coverslips were mounted on microscope slides (Thermo Fisher Scientific, Cat# 12-544-1) using Prolong Gold Antifade mounting media (Thermo Fisher Scientific, Cat# P10144). Coverslips were imaged using a Lionheart FX Automated Microscope (BioTek, Winooski, VT), taking images using the GFP, Texas Red, and DAPI filter cubes at each imaged position. For each condition, 20 images were collected and all cells in each image were analyzed using CellProfiler 3.0.0 (McQuin et al., 2018) (Broad Institute, Cambridge, MA). A modified CellProfiler analysis pipeline was created based on the example “Fruit fly cells” pipeline available at https://cellprofiler.org/examples/, excluding all cells whose cytoplasm was not contained fully within the image field. Calculated per-cell mean fluorescence intensity values were extracted for the GFP and Texas Red channels from each image and plotted in Prism.

### Chemical Library Screening

To screen for small molecule modulators of Pim-induced death, A549^N^ cells were seeded in two parallel 384-well plates. A library of 261 bioactive compounds (Selleck Chemicals, Cat# L2000) was diluted in SYTOX Green-containing DMEM from 2 mM to 50 μM in a dilution plate using a Versette liquid handler (Thermo Fisher Scientific). The day after seeding, the liquid handler was used to pipette SYTOX Green-containing media with either Pim or DMSO vehicle control into the assay plates containing A549^N^ cells. The compound library from the dilution plate was subsequently added to each assay plate, diluting the library 10X. The resulting assay plates contained A549^N^ cells treated with either 5 μM Pim or DMSO, plus the library compounds at a 5 μM dose. The plates were subsequently analyzed by STACK, imaging on an IncuCyte (Essen BioScience) every 4 h for 120 h total and processed and analyzed as described above. Three independent replicates of the library screen were conducted, with subsequent analysis performed using the mean of the lethal fraction (LF) for the three replicates at each timepoint.

Compound interactions were scored as follows. Normalized area under the curve (nAUC) metrics were calculated and compared using a Bliss model of drug independence. nAUC was calculated by generating an AUC value from the LF curve and dividing this value by the maximum possible AUC: an LF value of 1 at every timepoint measured. A mean nAUC value for Pim was calculated from DMSO control wells in the Pim-treated plates from all three replicates. Using the nAUC values for Pim (P) and each library compound (L) in the DMSO-treated plate, an expected nAUC (E) was calculated for each combination of Pim plus the library compound based on the Bliss model: E = P + L – (P * L). The deviation of the observed nAUC from this expected nAUC represented the enhancement or protection of Pim-induced death by the library compound.

### Quantification and Statistical Analysis

Lethal fraction scoring was performed using Microsoft Excel 14.6.0 (Microsoft Corporation, Redmond, WA). LED curve fitting was performed using Prism 8 (GraphPad Software, La Jolla, CA). Flow cytometry data were analyzed using FlowJo 10.6.1 (FlowJo LLC, Ashland, OR). Immunofluorescence images were quantified using CellProfiler 3.0.0 (Broad Institute, Cambridge, MA). Graphing and statistical analyses were performed using Prism 8. Figures were assembled using Adobe Illustrator (Adobe Systems, San Jose, CA). Statistical details of experiments and statistical tests used can be found in the main text, figure legends, and STAR Methods.

### Data and Code Availability

RNA-seq data are available in GEO under accession number GSE140172.

---

**Supplemental Figure 1, related to Figure 1.**
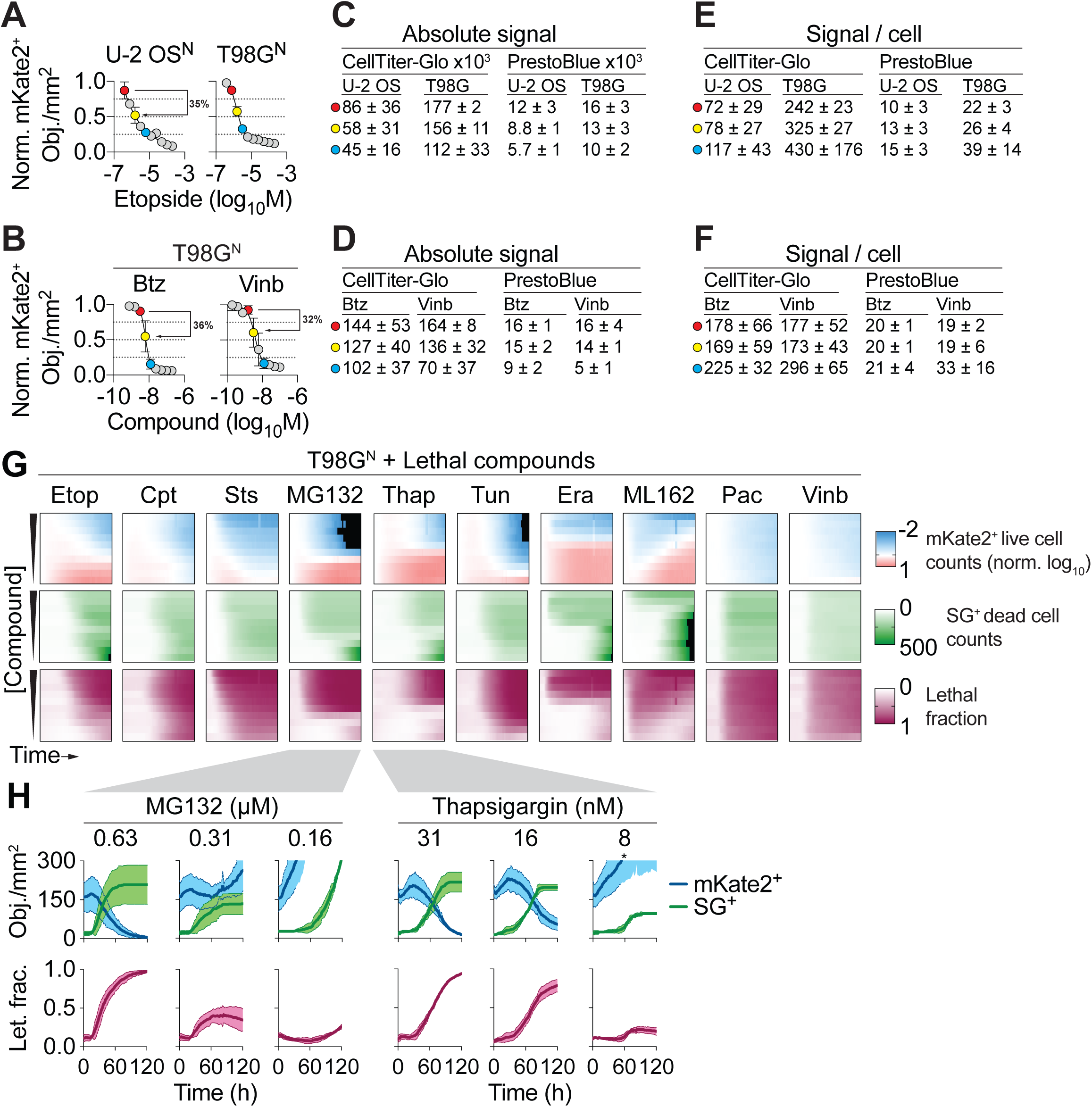
High-throughput assessment of fractional killing between conditions. (A,B) Assessment of live cell counts (mKate2 positive [mKate2^+^] objects) over lethal compound concentration-response series in two cell lines. Measurements were performed at 72 h. Btz: bortezomib, Vinb: vinblastine. Percent reduction in live cell counts between different compound concentrations is indicated by black lines and arrows and discussed in the main text. (C,D). Absolute metabolic viability measurements at select dose points. Colored circles in C and D correspond to specific doses indicated on the curves in A and B, respectively. (E,F). Metabolic viability measurements normalized per cell. Colored circles in E and F correspond to specific doses indicated on the curves in A and B, respectively. (G) Heatmap summaries of normalized, log_10_-scaled live mKate2 positive (mKate2+) live cell counts, SYTOX Green positive (SG+) dead cell counts, and integrated lethal fraction (Let. Frac.) scores over time (x-axis) and concentration (y-axis) for ten compounds in T98G^N^ cells. For the mKate2^+^ heatmaps, live cell counts were normalized to 1 at time = 0 and expressed on a log_10_ scale. Values more than 10-fold lower than the starting values are colored black. Etop: etoposide, Cpt: camptothecin, Sts: staurosporine, Thap: thapsigargin, Tun: tunicamycin, Era: erastin, Pac, paclitaxel. (H) Live and dead cell counts for T98G^N^ cells, extracted from the data shown in G. Data are from three independent experiments and represent mean ± SD (A-F, H) or mean (G).

**Supplemental Figure 2, related to Figure 2.**
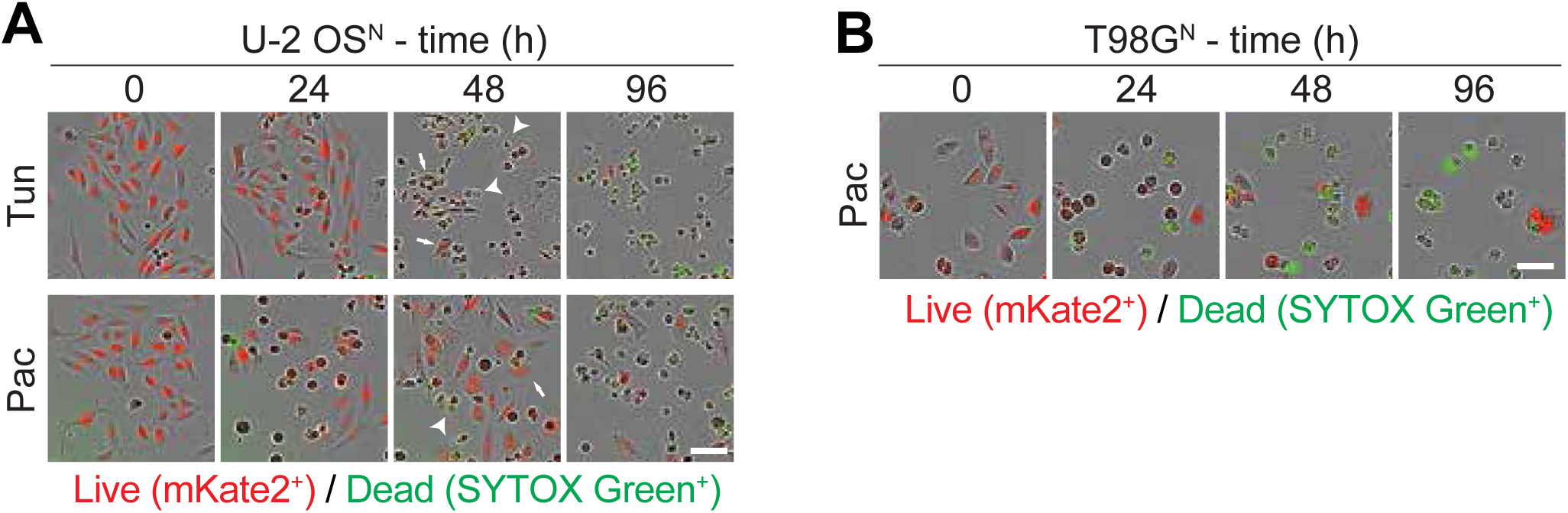
Kinetic analysis of FK. (A,B) Representative images for U-2 OS^N^ (A) and T98G^N^ (B) cells treated with the indicated lethal compounds over time. Arrows indicate example live cells, Arrowheads example dead cells. Images correspond to the underlying data analyzed as part of Figure 1. Scale bar = 75 µM. Tun: tunicamycin, Pac: paclitaxel.

**Figure S3, related to Figure 3.**
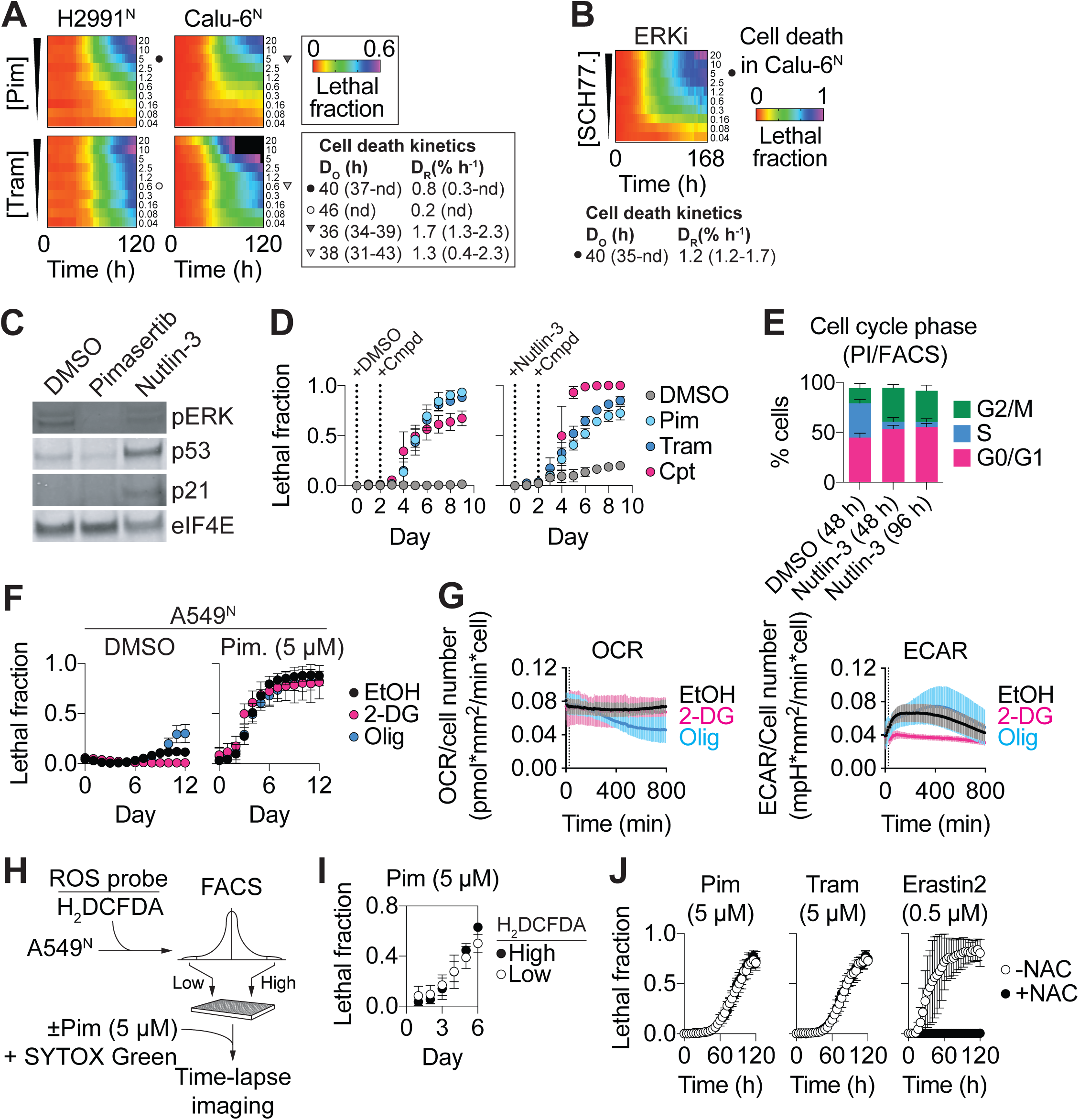
Investigating the MEKi-induced fractional killing. (A) Cell death summarized over time and MEKi concentration. Data are mean values from three independent experiments. Cell death kinetic parameters for select concentrations indicated by the symbols are shown at right, with 95% confidence intervals in brackets. nd: not determined. (B) Cell death summarized over time and SCH772984 concentration. Data are mean values from three independent experiments. Cell death kinetic parameters for one concentration indicated by the black circle are shown, with 95% confidence intervals in brackets. nd: not determined. (C) Phospho-ERK1/2 (pERK), p53 and p21 levels in A549^N^ cells ± nutlin-3 (10 µM, 48 h). (D) Cell death over time following pretreatment (2 day) ± nutlin-3 (10 µM). +Cmpd indicates the time of pimasertib (Pim, 5 µM), trametinib (Tram, 5 µM) or camptothecin (Cpt, 10 µM) addition. p53 stabilization has no effect on the kinetics of MEKi-induced cell death. (E) Analysis of cell cycle phase by PI/FACS ± nutlin-3 (10 µM) for the indicated times. (F) Cell death ± pimasertib ± vehicle (ethanol, EtOH), 2-deoxyglucose (2-DG, 10 mM) or oligomycin (Olig, 100 nM). Disruption of bioenergetics has no effect on FK. (G) Oxygen consumption (OCR) and extracellular acidification (ECAR) determined using a Seahorse assay in A549^N^ cells ± 2-DG or Olig as in F. (H) Outline of FACS-based scheme to isolate cells with high or low intracellular ROS as assessed using H_2_DCFDA. (I) Cell death over time for isolated H_2_DCFDA^High^ and H_2_DCFDA^Low^ sub-populations exposed to pimasertib (5 µM). (J) Cell death over time in A549^N^ cells ± MEKis ± N-acetylcysteine (NAC, 1 mM). Erastin2 is a positive control compound that induces oxidative cell death. Data in D-G, I and J are mean ± SD from at least three independent experiments.

**Figure S4, related to Figure 4.**
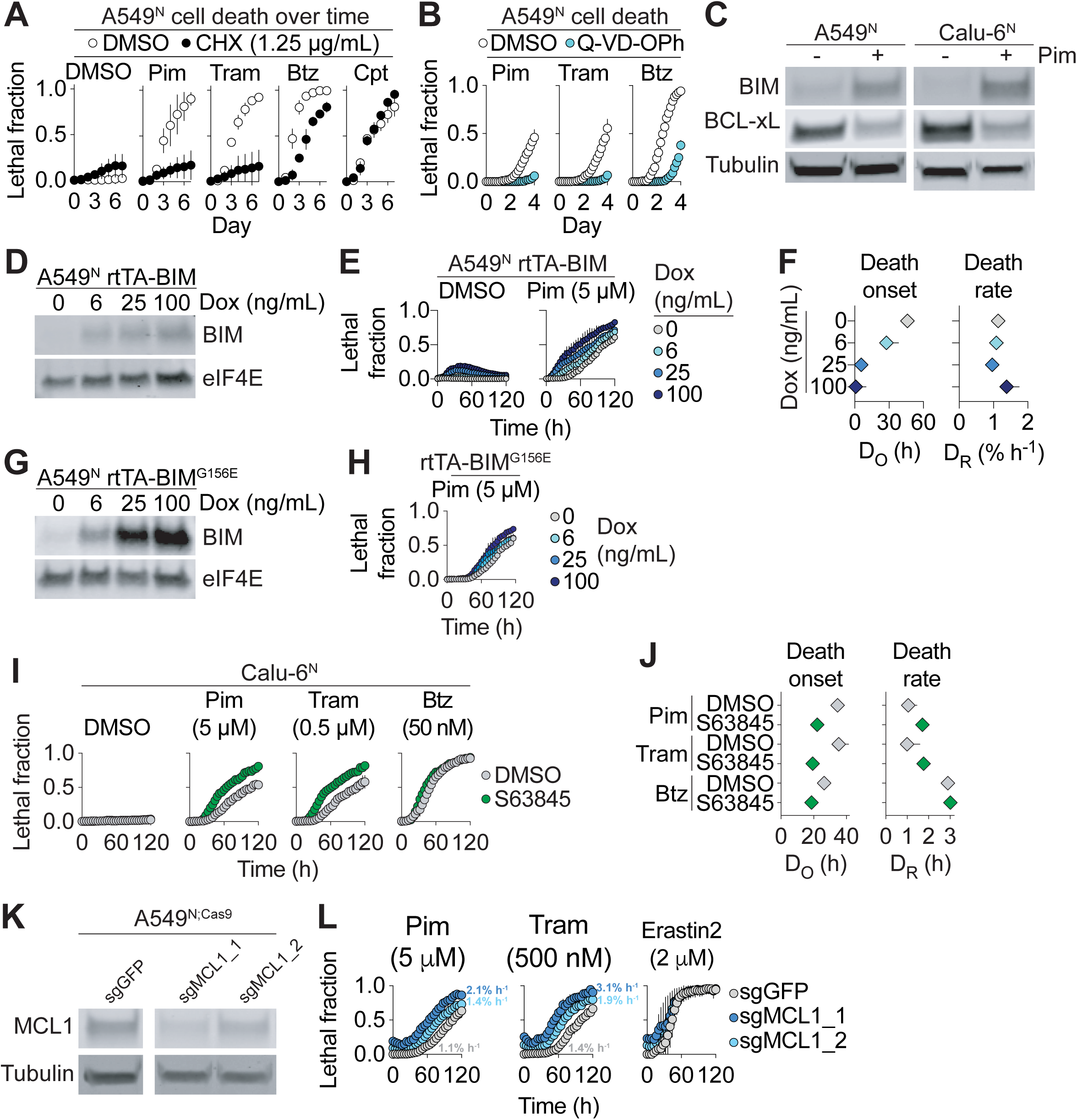
(A) Cell death in response to treatment with pimasertib (Pim, 5 µM), trametinib (Tram, 5 µM), bortezomib (Btz, 50 nM), camptothecin (Cpt, 1 µM) ± cycloheximide (CHX). (B) Cell death ± Q-VD-OPh (50 µM). Pim, Tram and Btz treatments are as in A. (C) Protein expression ± pimasertib (5 µM, 48 h). (D) Expression of BIM in response to doxycycline (Dox). (E) Cell death over time ± Pim combined with Dox-inducible expression of wild-type BIM. (F) Cell death kinetic parameters computed from the data in E. (G) Expression of the inactive BIM^G156E^ mutant + Dox. (H) Cell death over time in response to Dox-inducible expression of mutant BIM^G156E^ + Pim. (I) Cell death in Calu-6^N^ cells in response to compound treatment ± the selective MCL1 inhibitor S63845 (5 µM). (J) Cell death kinetic parameters computed from the data in I. (K) Expression of MCL1 in A549^N;Cas9^ cells transduced with short guide RNAs against *GFP* or *MCL1*. (L) Cell death over time in the cells lines from K, treated as indicated. D_R_ values computed from each curve are indicated for MEKis. Data are from at least three independent experiments and represent mean ± SD (A,B,E,H,I,L) or mean ± 95% C.I (F,J).

**Figure S5, related to Figure 5.**
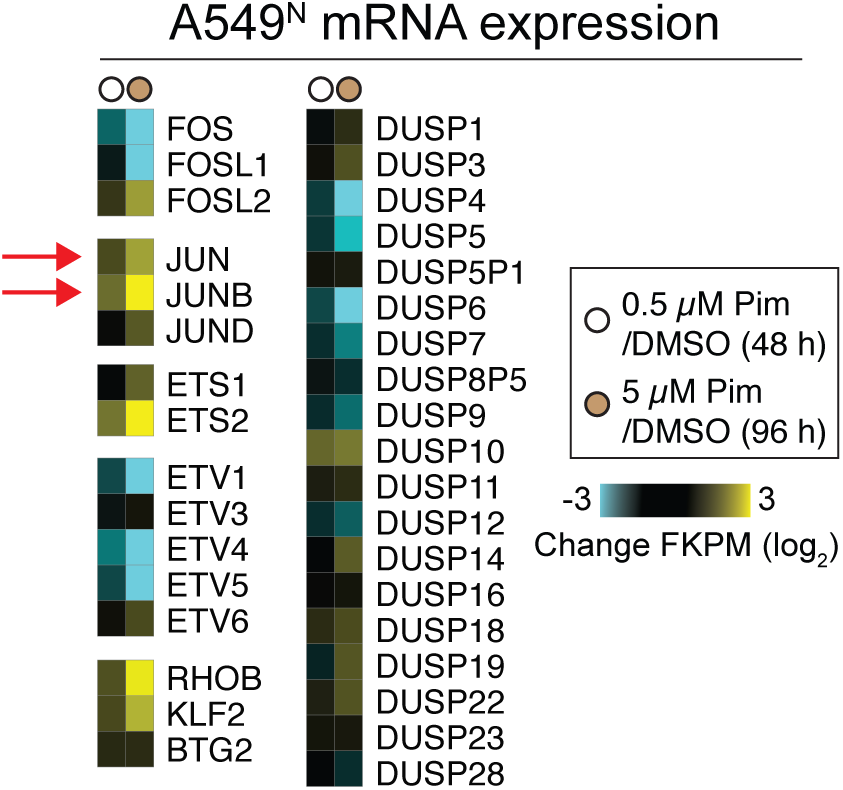
Subset of the RNA-seq data presented in Figure 4A, showing relative changes in immediate early genes mRNA. The heatmap represents mean values for two independent biological replicates.

KEY RESOURCES TABLE

KEY RESOURCES TABLE
| REAGENT or RESOURCE | SOURCE | IDENTIFIER |
| --- | --- | --- |
| **Antibodies** |  |  |
| Anti-Tubulin alpha (mouse) | Fisher Scientific | Cat# MS581P1 |
| Anti-eIF4E (mouse) | Santa Cruz Biotechnology | Cat# sc-9976 |
| Anti-p44/42 MAPK (Erk1/2) Rabbit mAb | Cell Signaling Technology | Cat# 4695 |
| Anti-phospho-p44/42 MAPK (Erk1/2) Rabbit mAb | Cell Signaling Technology | Cat# 4370 |
| Anti-DUSP4 (MKP2) Rabbit mAb | Cell Signaling Technology | Cat# 5149 |
| Anti-Bim Rabbit mAb | Cell Signaling Technology | Cat# 2933 |
| Anti-Bcl-xL Rabbit mAb | Cell Signaling Technology | Cat# 2764 |
| Anti-Mcl-1 Rabbit mAb | Cell Signaling Technology | Cat# 94296 |
| Anti-phospho-c-Jun (Ser73) Rabbit mAb | Cell Signaling Technology | Cat# 3270 |
| Anti-c-Jun Rabbit mAb | Cell Signaling Technology | Cat# 9165 |
| Anti-phospho-S6 Ribosomal Protein (Ser235/236) Rabbit mAb | Cell Signaling Technology | Cat# 4858 |
| Anti-S6 Ribosomal Protein Rabbit mAb | Cell Signaling Technology | Cat# 2217 |
| Anti-phospho-4E-BP1 (Thr37/46) Rabbit mAb | Cell Signaling Technology | Cat# 9459 |
| Anti-4E-BP Rabbit mAb | Cell Signaling Technology | Cat# 9644 |
| IRDye 800CW Donkey anti-Rabbit IgG | Licor | Cat# 926-32213 |
| IRDye 800CW Donkey anti-Mouse IgG | Licor | Cat# 926-32212 |
| IRDye 680LT Donkey anti-Rabbit IgG | Licor | Cat# 926-68023 |
| IRDye 680LT Donkey anti-Mouse IgG | Licor | Cat# 926-68022 |
| Goat polyclonal anti-rabbit highly cross-adsorbed secondary antibody, Alexa Fluor 568 | Life Technologies | Cat# A11036 |
| Goat polyclonal anti-mouse highly cross-adsorbed secondary antibody, Alexa Fluor 488 | Life Technologies | Cat# A11029 |
| **Bacterial and Virus Strains** |  |  |
| NucLight Red lentivirus reagent (EF1a, Puro) Nuclear-localized mKate2 (Nuc::mKate2) | Essen BioSciences | Cat# 4265 |
| NucLight Red lentivirus reagent (EF1a, Bleo) Nuclear-localized mKate2 (Nuc::mKate2) | Essen BioSciences | Cat# 4627 |
| **Biological Samples** |  |  |
| N/A |  |  |
| **Chemicals, Peptides, and Recombinant Proteins** |  |  |
| Puromycin | Life Technologies | Cat# A11138-03 |
| Geneticin | Thermo Fisher Scientific | Cat# 11811-031 |
| Blasticidin | Thermo Fisher Scientific | Cat# A1113902 |
| Camptothecin | Selleck Chemicals | Cat# S1288 |
| Vinblastine | Selleck Chemicals | Cat# S1248 |
| Pimasertib | Selleck Chemicals | Cat# S1457 |
| Trametinib | Selleck Chemicals | Cat# S2673 |
| SCH772984 | Selleck Chemicals | Cat# S7101 |
| ABT-737 | Selleck Chemicals | Cat# S1002 |
| A-1155463 | Selleck Chemicals | Cat# S7800 |
| Nutlin-3 | Selleck Chemicals | Cat# S1061 |
| S63845 | Cayman Chemical Company | Cat# 21131 |
| Staurosporine | ApexBio | Cat# A8192 |
| N-acetylcysteine | Millipore Sigma | Cat# A8199 |
| Thapsigargin | Millipore Sigma | Cat# T9033 |
| Tunicamycin | Millipore Sigma | Cat# T7765 |
| Paclitaxel | Millipore Sigma | Cat# T7191 |
| JNK Inhibitor VIII | Millipore Sigma | Cat# 420135 |
| 2-deoxyglucose | Millipore Sigma | Cat# D8375 |
| Oligomycin | Millipore Sigma | Cat# O4876 |
| Bortezomib | Thermo Fisher Scientific | Cat# NC0587961 |
| Etoposide | Thermo Fisher Scientific | Cat# ICN19391825 |
| MG-132 | Thermo Fisher Scientific | Cat# 17-485 |
| Q-VD-OPh | Thermo Fisher Scientific | Cat# OPH00101M |
| Erastin2 (35MEW28) | Dixon et al., 2014 | N/A |
| Bioactive chemical library (261 compounds) | Selleck Chemicals | Cat# L2000 |
| Cycloheximide | Sigma-Aldrich | Cat# C7698-1G |
| Polybrene | Sigma-Aldrich | Cat# H9268-5G |
| Sytox Green | Life Technologies | Cat# S7020 |
| PolyJet | SigmaGen Laboratories | Cat#SL100688 |
| H<sub>2</sub>DCFDA | Thermo Fisher Scientific | Cat# D-399 |
| DAPI (4',6-Diamidino-2-Phenylindole, Dihydrochloride) | Thermo Fisher Scientific | Cat# D1306 |
| FastDigest BstXI | Thermo Fisher Scientific | Cat# FD1024 |
| FastDigest Bpu1120I | Thermo Fisher Scientific | Cat# FD0094 |
| **Critical Commercial Assays** |  |  |
| Gateway LR Clonase II Enzyme Mix | Thermo Fisher | Cat# 11791-020 |
| Gibson Assembly Master Mix | New England Biolabs | Cat# E2611S |
| QIAshredder RNA Extraction Column Kit | Qiagen | Cat# 79654 |
| RNeasy Plus RNA Extraction Kit | Qiagen | Cat# 74134 |
| TaqMan Reverse Transcriptase Kit | TaqMan | Cat# N8080234 |
| QIAprep Spin Miniprep Kit | Qiagen | Cat# 27106 |
| QIAquick PCR Purification Kit | Qiagen | Cat# 28106 |
| Q5 Site-Directed Mutagenesis Kit | New England Biolabs | Cat# E0554S |
| SYBR Green Master Mix | Life Technologies | Cat# 4367659 |
| BCA Protein Assay Kit | Thermo Fisher | Cat# 23225 |
| CellTiter-Glo Luminescent Cell Viability Assay | Promega | Cat# G7570 |
| PrestoBlue Cell Viability Reagent | Thermo Fisher | Cat# A13261 |
| **Deposited Data** |  |  |
| N/A |  |  |
| **Experimental Models: Cell Lines** |  |  |
| U-2 OS | ATCC | HTB-96 |
| U-2 OS<sup>N</sup> | Forcina et al., 2017 | N/A |
| T98G | ATCC | CRL-1960 |
| T98G<sup>N</sup> | Forcina et al., 2017 | N/A |
| A549 | ATCC | CCL-185 |
| A549<sup>N</sup> | Forcina et al., 2017 | N/A |
| A549<sup>N</sup>; KTR | This paper | N/A |
| A549<sup>N</sup>; rtTA-BCL-xL | This paper | N/A |
| A549<sup>N</sup>; rtTA-BIM | This paper | N/A |
| A549<sup>N</sup>; rtTA-BIM-G156E | This paper | N/A |
| A549<sup>N</sup>; CMV-Empty | Cao et al., 2019 | N/A |
| A549<sup>N</sup>; CMV-MCL1 | This paper | N/A |
| A549<sup>N</sup>; Cas9 | This paper | N/A |
| A549<sup>N</sup>; Cas9; sgGFP | This paper | N/A |
| A549<sup>N</sup>; Cas9; sgMCL1 #1 | This paper | N/A |
| A549<sup>N</sup>; Cas9; sgMCL1 #2 | This paper | N/A |
| A549<sup>N</sup>; CRISPR Control | This paper | N/A |
| A549<sup>N</sup>; MCL1-LOF1 | This paper | N/A |
| A549<sup>N</sup>; MCL1-LOF2 | This paper | N/A |
| Calu-6 | ATCC | HTB-56 |
| Calu-6<sup>N</sup> | This paper | N/A |
| HEK293T | ATCC | CRL-3216 |
| **Experimental Models: Organisms/Strains** |  |  |
| N/A |  |  |
| **Oligonucleotides** |  |  |
| See Table S2. |  |  |
| **Recombinant DNA** |  |  |
| pLenti CMVTRE3G BCL2L1 Neo | This paper | N/A |
| pLenti CMVTRE3G BCL2L11 Neo | This paper | N/A |
| pLenti CMVTRE3G BCL2L11 G156E Neo | This paper | N/A |
| pLenti CMV MCL1 Puro | This paper | N/A |
| pMCB306 MCL1 sgRNA #1 | This paper | N/A |
| pMCB306 MCL1 sgRNA #2 | This paper | N/A |
| **Other Plasmids** |  |  |
| pLentiCMV Puro DEST ERKKTRClover | Addgene | Cat# 59150 |
| pLentiPGK Puro DEST p38KTRClover | Addgene | Cat# 59152 |
| pLenti CMV rtTA Blast (w756-1) | Addgene | Cat# 26429 |
| pLenti CMVTRE3G eGFP Neo (w821-1) | Addgene | Cat# 27569 |
| pLenti CMV Puro DEST (w118-1) | Addgene | Cat# 17452 |
| pMCB306 | Addgene | Cat# 89360 |
| lentiCas9-Blast | Addgene | Cat# 52962 |
| **Software and Algorithms** |  |  |
| Prism | GraphPad Software | N/A |
| Excel | Microsoft Corp. | N/A |
| FlowJo | FlowJo LLC. | N/A |
| CellProfiler | Broad Institute | N/A |
| **Data availability** |  |  |
| N/A |  |  |

**Table S1.**
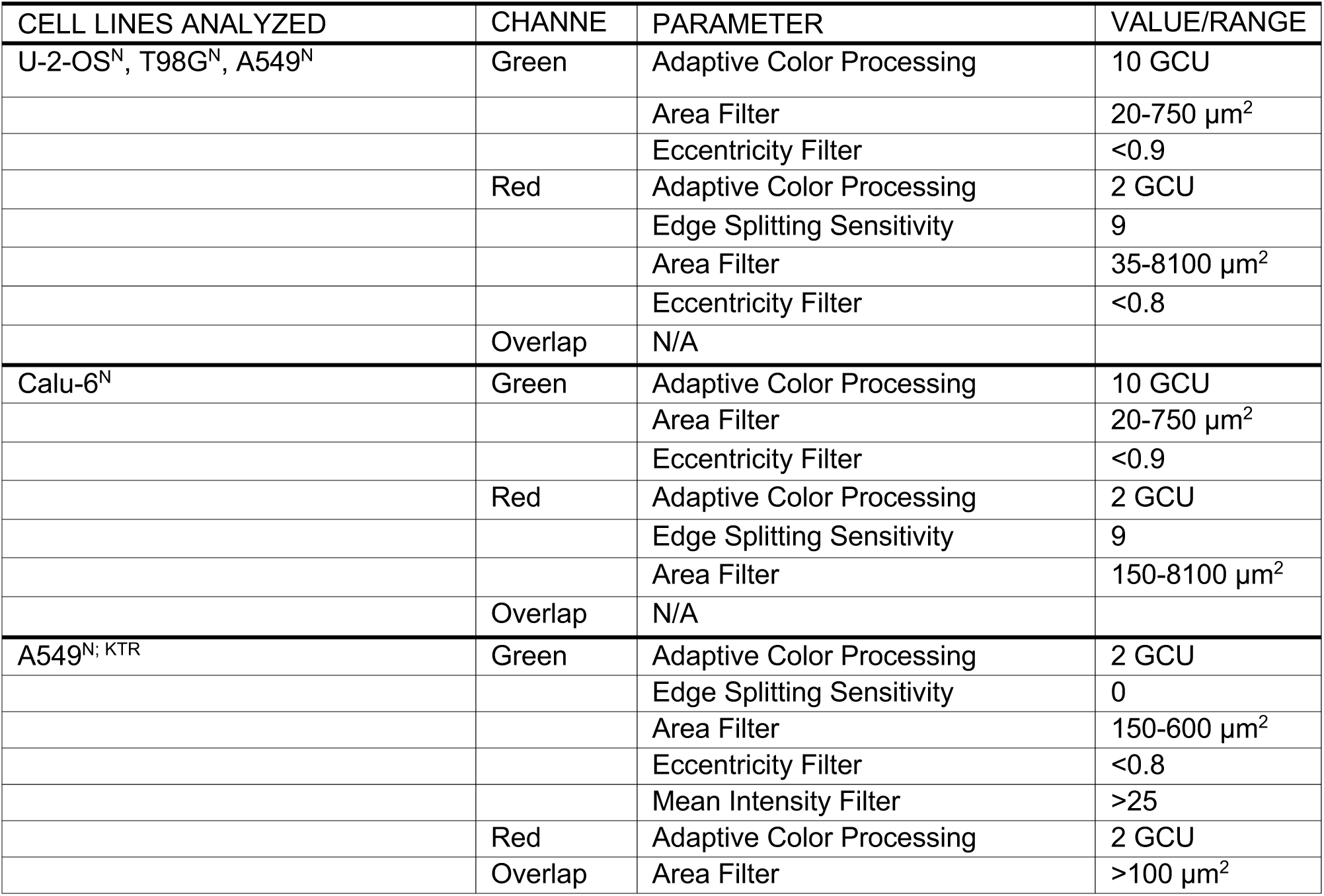
Related to STAR Methods. Parameters for IncuCyte analysis.

**Table S2.**
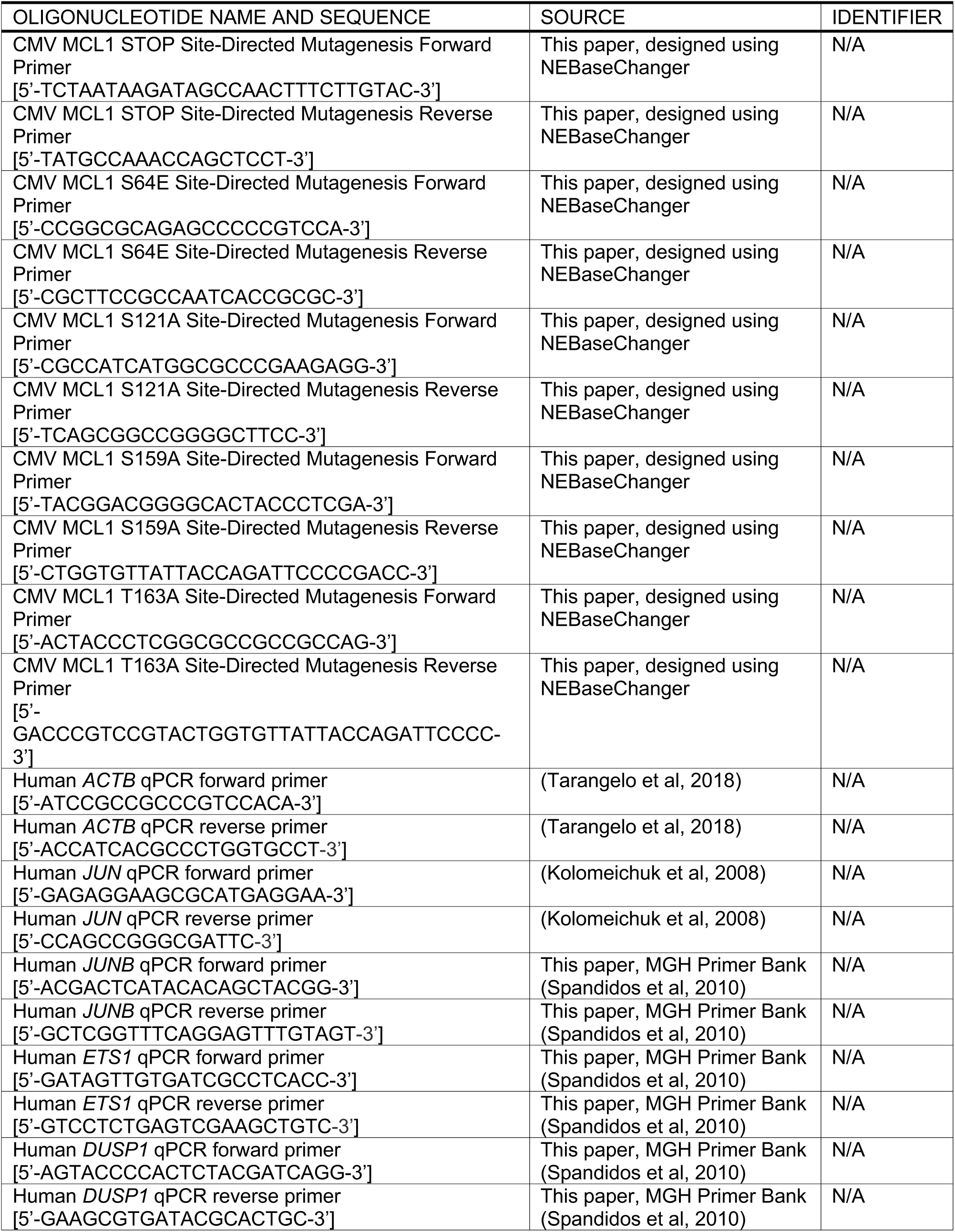
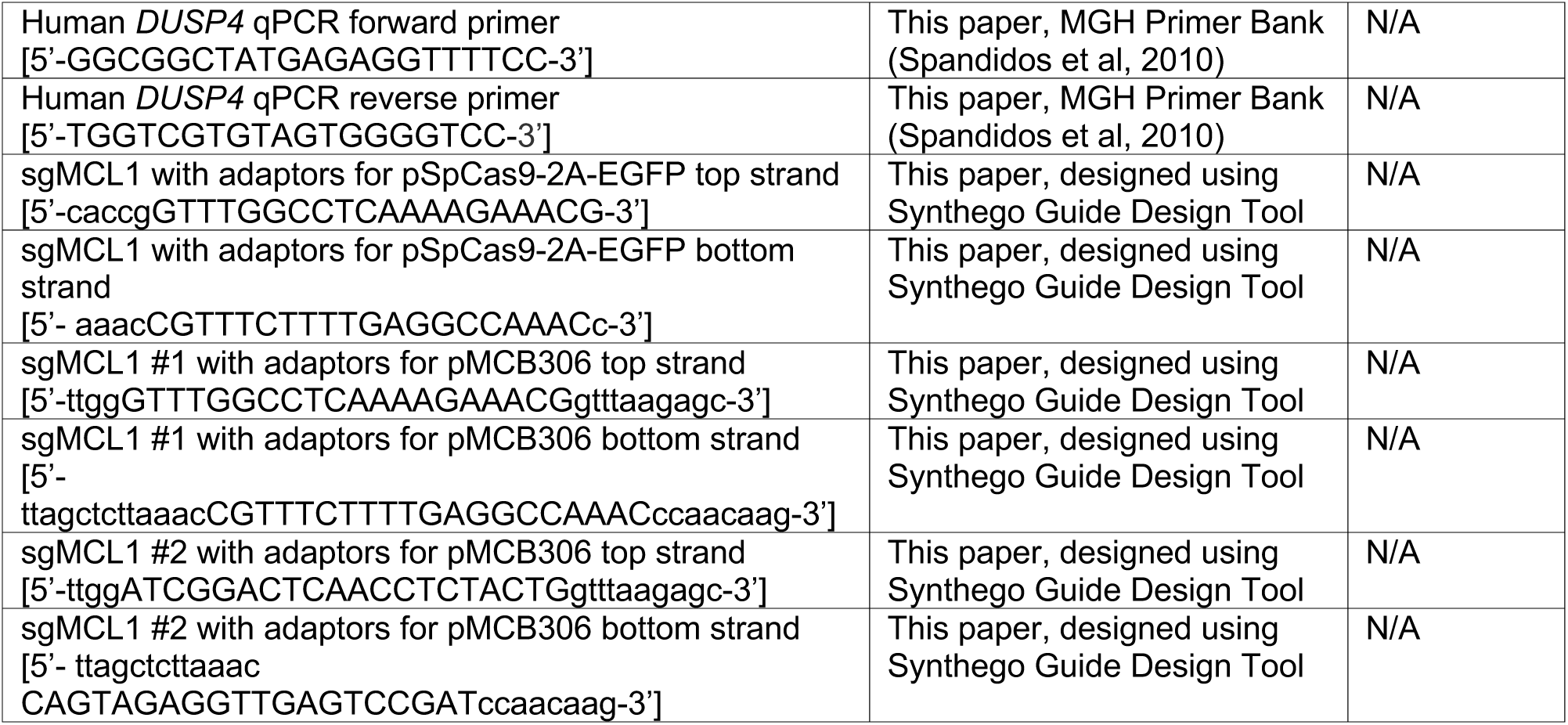
Related to STAR Methods. Oligonucleotides used in this study.

